# The *Silene latifolia* genome and its giant Y chromosome

**DOI:** 10.1101/2023.09.21.558754

**Authors:** Carol Moraga, Catarina Branco, Quentin Rougemont, Paris Veltsos, Pavel Jedlička, Aline Muyle, Melissa Hanique, Eric Tannier, Xiaodong Liu, Eddy Mendoza-Galindo, Claire Lemaitre, Peter D. Fields, Corinne Cruaud, Karine Labadie, Caroline Belser, Jerome Briolay, Sylvain Santoni, Radim Cegan, Raquel Linheiro, Ricardo C. Rodríguez de la Vega, Gabriele Adam, Adil El Filali, Vinciane Mossion, Adnane Boualem, Raquel Tavares, Amine Chebbi, Richard Cordaux, Cécile Fruchard, Djivan Prentout, Amandine Velt, Bruno Spataro, Stephane Delmotte, Laura Weingartner, Helena Toegelová, Zuzana Tulpová, Petr Cápal, Hana Šimková, Helena Štorchová, Manuela Krüger, Oushadee A. J. Abeyawardana, Douglas R. Taylor, Matthew S. Olson, Daniel B. Sloan, Sophie Karrenberg, Lynda F. Delph, Deborah Charlesworth, Tatiana Giraud, Abdelhafid Bendahmane, Alex Di Genova, Amin Madoui, Roman Hobza, Gabriel A. B. Marais

## Abstract

In some species, the Y is a tiny chromosome but the dioecious plant *Silene latifolia* has a giant ∼550 Mb Y chromosome, which has remained unsequenced so far. Here we used a hybrid approach to obtain a high-quality male *S. latifolia* genome. Using mutants for sexual phenotype, we identified candidate sex-determining genes on the Y. Comparative analysis of the sex chromosomes with outgroups showed the Y is surprisingly rearranged and degenerated for a ∼11 MY-old system. Recombination suppression between X and Y extended in a stepwise process, and triggered a massive accumulation of repeats on the Y, as well as in the non-recombining pericentromeric region of the X, leading to giant sex chromosomes.

**One-Sentence Summary:** This work uncovers the structure, function, and evolution of one of the largest giant Y chromosomes, that of the model plant *Silene latifolia*, which is almost 10 times larger than the human Y, despite similar genome sizes.

## Main Text

Among the multiple paths the evolution of sex chromosomes can take, some lead to giant Y chromosomes (*1,2*). Giant Y chromosomes have been first identified in dioecious plant species (*3*) but also exist in animals (*4*). In the last decade, great advances have been made in studying plant sex chromosomes thanks to genomics and bioinformatics (*5,6*), but no giant plant Y chromosome has been assembled yet. Giant Y chromosomes may result from massive accumulation of transposable elements (TEs) but their chromosome organization, their precise role in sex determination, and their evolution remain poorly known (*3*).

*Silene latifolia* (Caryophyllaceae) is a dioecious plant that has been studied since Darwin’s time (*7*). *S. latifolia* has an XY sex-determination system, which was discovered 100 years ago (*8*). The Y is ∼550 Mb, and the X ∼400 Mb, of the total haploid genome size of ∼2.7 Gb (*9*). Genetic maps show that the X and the Y are largely non-recombining and share only a single pseudo-autosomal region (PAR) (*10*). Recombination has been suppressed progressively forming groups of X/Y gene pairs with differing synonymous divergence, called evolutionary strata (*11,12*). The repeat-richness (*13*) and size of the *S. latifolia* Y have, however, so far prevented its assembly. Mutants with deletions on the Y chromosome and altered sex phenotypes indicate the presence of three sex-determining regions (*14*): one female-suppressing region (carrying a gynoecium-suppressing factor, GSF) and two male-promoting (carrying a stamen-promoting factor - SPF and male-fertiliy factor - MFF) (*15*). A GSF candidate gene has recently been proposed (*16*), but the other sex-determining genes remain unknown.

Here we used an Oxford-Nanopore-Technology (ONT)-based sequencing approach to obtain the *S. latifolia* genome, in order to study the repeat-rich Y chromosome. We also used high-quality genome assemblies of closely related non-dioecious *Silene* species as outgroups to make inferences about the evolution of the *S. latifolia* sex chromosomes. We sequenced mutants with Y deletions for three sexual phenotypes (hermaphrodites and asexuals with early/intermediate and late anther development arrest) in order to pinpoint candidate sex-determining genes/regions. Furthermore, we generated expression data at two critical stages for male and female flower development to help identify sex-determining genes.

### The structure and gene content of the sex chromosomes

To assemble the complex *S. latifolia* genome, we produced an inbred population from 17 generations of brother-sister crosses to reduce heterozygosity (see Material and Methods for the details of the sequencing, assembly and annotation methodologies). We selected one male and one female individual from this inbred population for sequencing. The male was sequenced using ONT at 100X coverage to ensure 50X coverage for both sex chromosomes (Table S1). Assembly was performed using an initial hybrid long-read approach including Flye (Figure S1). The ONT contigs were polished using 200X of short Illumina reads and then scaffolded using Bionano optical mapping data followed by Omni-C contact data (see Table 1 for genome statistics and Table S2 for more details). The polished consensus sequence reached a phred quality score of ∼30 (1/1000 base error). All scaffolds were anchored onto our genetic map, except scaffolds 1, 12, 13, 16 and 41 (Figure S2). These unplaced scaffolds were classified as likely fragments of the Y chromosome based on female/male sequence depths and mapping of previously well-characterized sex-linked genes (see below). They were assembled into a single sequence. Re-mapping the Omni-C data onto the chromosome sequences obtained in this manner further improved the scaffolding of the X and Y chromosomes. The final chromosome sequences show high consistency with the contact data (Figure S3). During the assembly process, the two haplotypes of each sequenced region were collapsed into a single sequence. We present this a haploid version of the *S. latifolia* genome with a single pseudomolecule for each chromosome, including separate assemblies of X and the Y chromosomes, except for the PARs on the X and Y, which were also collapsed (the resulting single PAR sequence is assembled on the X chromosome only (see below). The sizes of the genome and sex chromosomes obtained closely match the previous C-value-based size estimates (Table 1). Gene annotation identified 35,436 protein-coding genes. BUSCO scores were high at different steps of the assembly/annotation process and the final score was 86.5% (Tables 1 and S2). Repeat analysis revealed that 79.24% of the

**Table 1.**
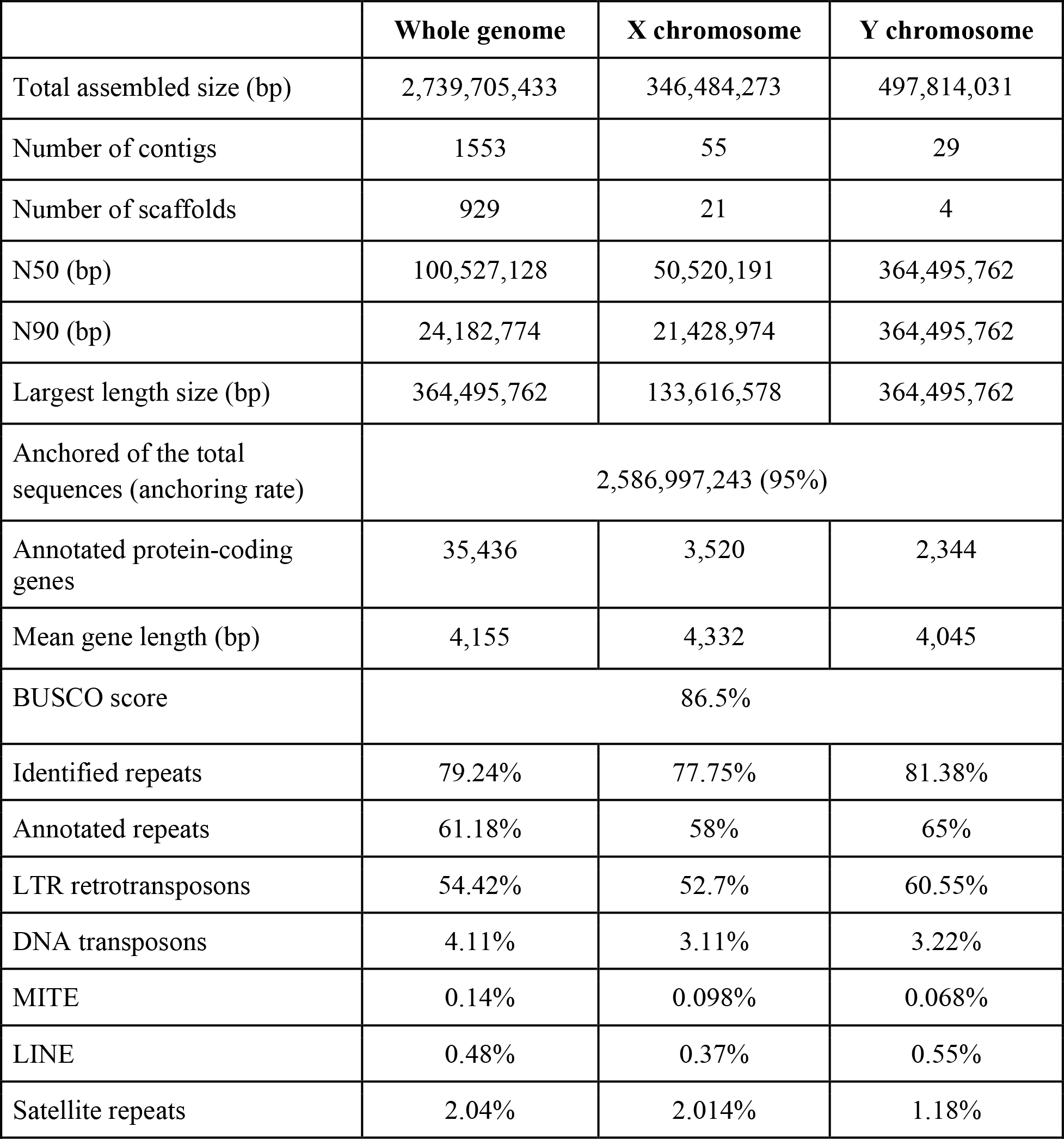
Statistics for the genome and the sex chromosomes. All assembly metrics were computed by the total assembly and sex chromosomes.

|  | **Whole genome** | **X chromosome** | **Y chromosome** |
| --- | --- | --- | --- |
| Total assembled size (bp) | 2,739,705,433 | 346,484,273 | 497,814,031 |
| Number of contigs | 1553 | 55 | 29 |
| Number of scaffolds | 929 | 21 | 4 |
| N50 (bp) | 100,527,128 | 50,520,191 | 364,495,762 |
| N90 (bp) | 24,182,774 | 21,428,974 | 364,495,762 |
| Largest length size (bp) | 364,495,762 | 133,616,578 | 364,495,762 |
| Anchored of the total sequences (anchoring rate) | 2,586,997,243 (95%) |  |  |
| Annotated protein-coding genes | 35,436 | 3,520 | 2,344 |
| Mean gene length (bp) | 4,155 | 4,332 | 4,045 |
| BUSCO score | 86.5% |  |  |
| Identified repeats | 79.24% | 77.75% | 81.38% |
| Annotated repeats | 61.18% | 58% | 65% |
| LTR retrotransposons | 54.42% | 52.7% | 60.55% |
| DNA transposons | 4.11% | 3.11% | 3.22% |
| MITE | 0.14% | 0.098% | 0.068% |
| LINE | 0.48% | 0.37% | 0.55% |
| Satellite repeats | 2.04% | 2.014% | 1.18% |

*S. latifolia* male genome consists of repeats (Table S3). Annotated repeats represented 61.18% of the *S. latifolia* genome with a very high abundance of two LTR retrotransposons *Copia* (20.73%) and *Gypsy* (33.69%). Telomere-associated repeats were found at the expected locations for all chromosome sequences, which constitute telomere-to-telomere assemblies (Figure 1A). Genes are concentrated in the chromosomal arms and are sparse in the pericentromeric regions, and vice versa for repeats, as in many eukaryotic genomes (*17*).

**Fig. 1.**
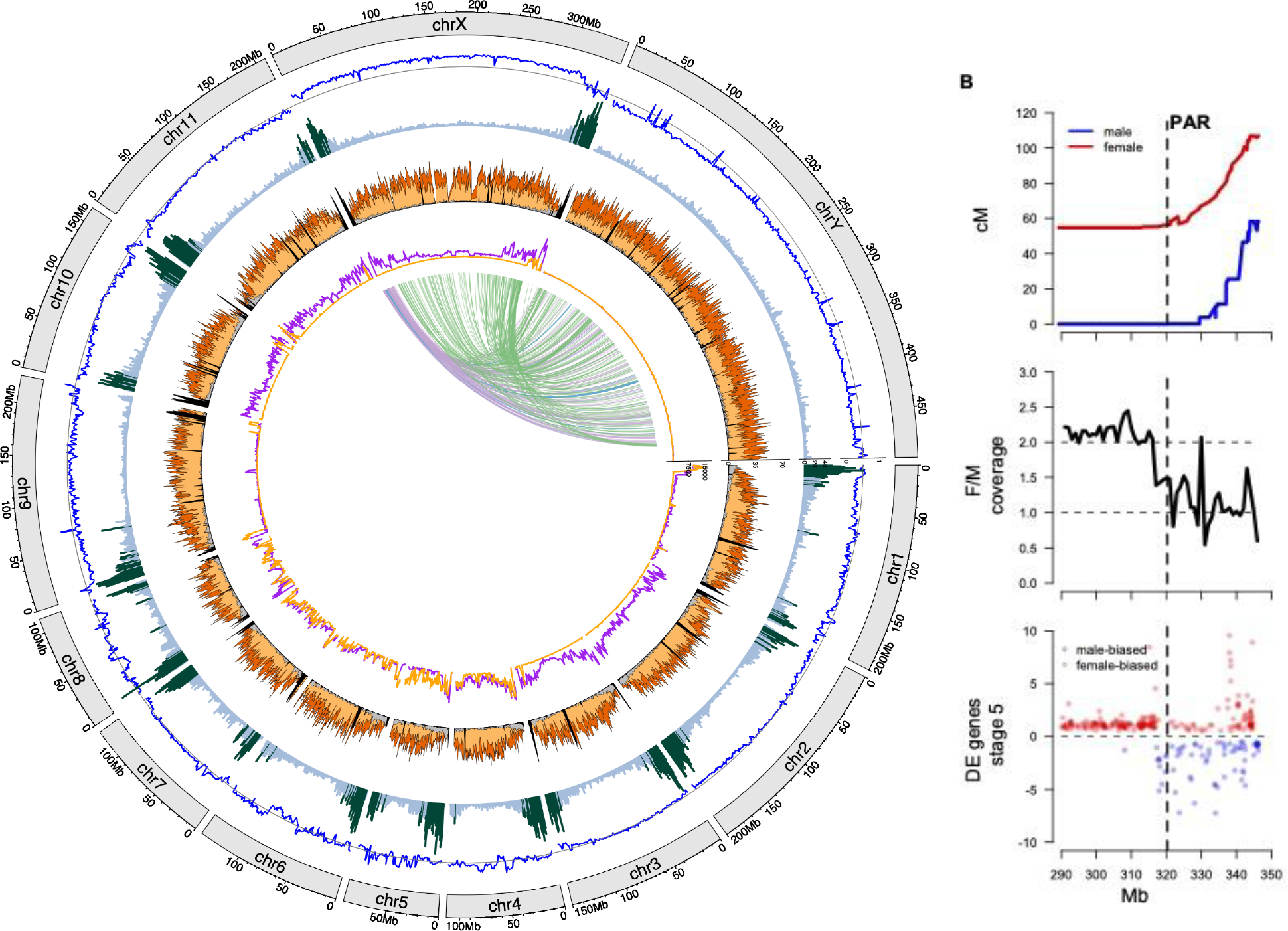
View of the assembly of the *S. latifolia* male genome assembly. (A) Circos plot of the *S. latifolia* male genome. Circles included in the Circos plot are from outer to inner circles: 1. The coverage Female/Male ratio distribution. 2. Gene density highlights the most populated regions (dark green); they are defined as those with a density larger than the average plus 1 sd. 3. Family repeats density distribution showing satellite elements (black), LTR elements: Ty3/Gypsy (orange), Ty1/Copia (yellow), LINE (violet), and Helitron (grey). 4. SNP Female and male distribution (purple for female and orange for male). This analysis is consistent with the sequenced male being highly homozygous, although some chromosomes show heterozygosity. It is also consistent with the female being a sister of the male. 5. Rearrangements between X and Y chromosomes based on the gametologs. (B) Zoom in on the X chromosome showing recombination in males (blue) and females (red) defining the pseudo-autosomal boundary at 321 Mb, female/male sequence coverage ratio, and significant differential expression between male and female flowers (stage 5); the full analysis of differential gene expression is shown in Figure S11. All panels have data summarised in 1Mb windows.

Our assemblies of the X and the Y chromosomes are of high quality. In systems in which the X and Y are differentiated (with SNPs and indels) and using stringent mapping parameters, female/male sequence-depth ratio of ∼1, ∼2 and ∼0 for autosomes, X, and Y chromosomes, respectively, are expected, and were observed in our data (Figure 1A). A smaller set of experimentally validated sex-linked genes (compiled in Muyle et al. 2018) also mapped as expected to their previously assigned X or Y position (Figures 2 and 3). The X chromosome sequence obtained is 346 Mb long. The distribution of genes and repeats (in particular centromere- and telomere associated repeats) is also as expected for a typical metacentric chromosome (Figure 1A). Sex-specific recombination data and other data identified the pseudo-autosomal region (Figure 1B). The PAR is a small gene-rich region (25 Mb with 1,286 genes). The Y chromosome assembly is 497 Mb long (522 Mb if the PAR is included) and includes some of the largest scaffolds (1 - 364.5 Mb, 12 - 45.7 Mb, 13 - 41.9 Mb).

**Fig. 2.**
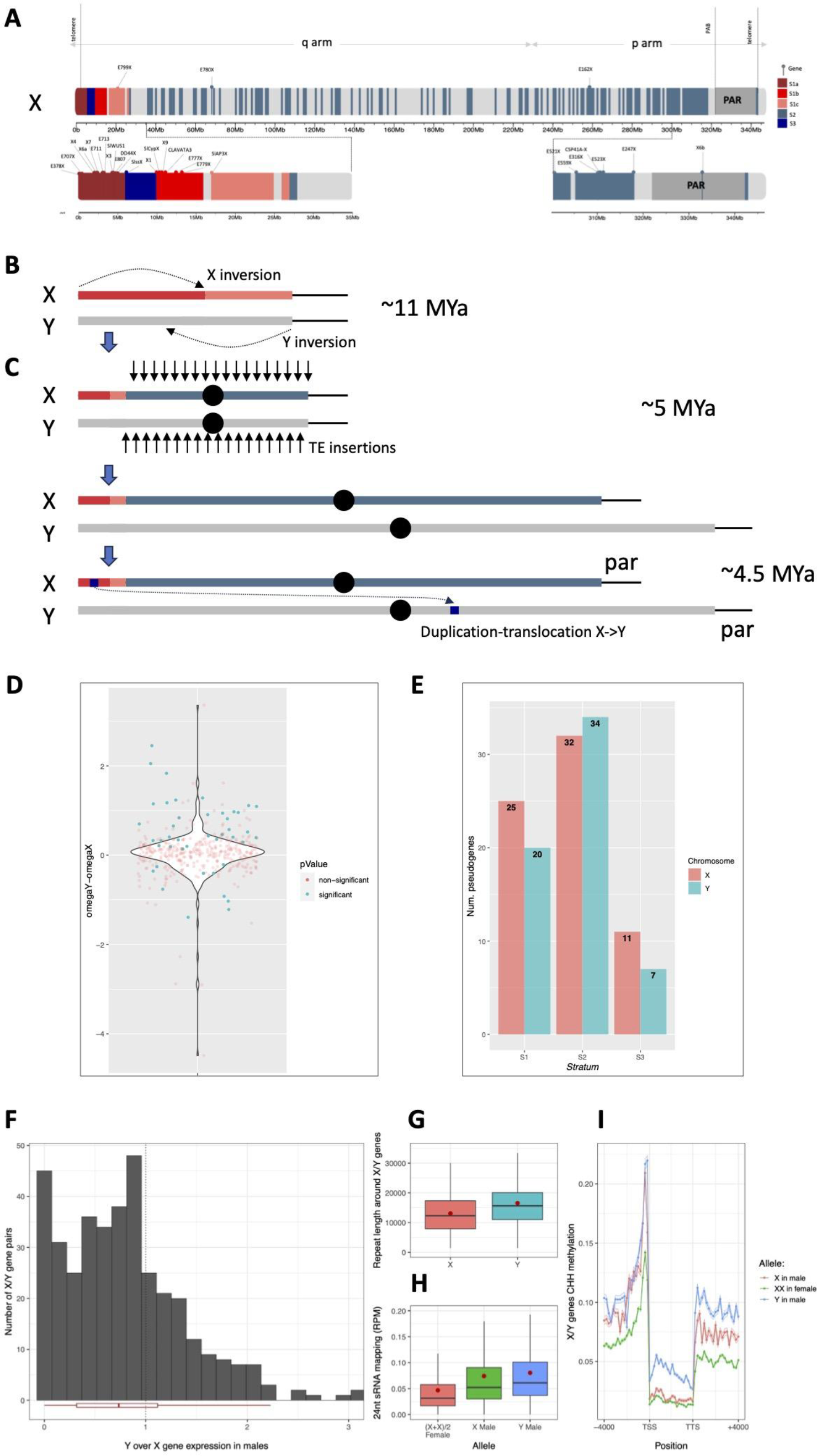
The evolution of the sex chromosomes. (A) The X chromosome sequence. The 417 gametologs genes are placed and colored according to their evolutionary stratum (S1a, S1b, S1c, S2 and S3). Zoom in of the extremities of the X chromosome are also shown. Genes previously characterized in the literature are identified by a ball-headed pin colored according to their stratum and located over the X chromosome sequence. (B) Schematic view of the first step of the evolution of the sex chromosomes with a zoom in on the first 35 Mb of the p arm of the X chromosome. The formation of stratum 1 (from S1a to S1c) is shown. Strata are not represented on the Y. (C) Global view of the other steps of the evolution of the sex chromosomes (full chromosomes are shown). (D) Violin plot of dN/dS differences among X and Y copies, gametologs pairs with significant differences (p-value < 0.05) are shown in blue. (E) Genes with premature stop codons on the sex chromosomes shown by strata. (F) Distribution of Y over X gene expression ratio in four *S. latifolia* males in flower buds. The boxplots below the distribution represent the median, the first and third quartiles (hinges) and 1.5 times the distance between the first and third quartiles (whiskers). (G) Sum of repeat lengths around X/Y gene pairs, from 4000 bp upstream to 4000 bp downstream of the gene. If repeats went beyond the borders (+/-4000bp), their length was not included. (H) Mapping of 24nt small RNA (in reads per million mapped reads, abbreviated RPM) on X/Y gene pairs for three *S. latifolia* females and three males in flower buds and leaves. The boxplots represent the median, the first and third quartiles (hinges) and 1.5 times the distance between the first and third quartiles (whiskers). The red dot stands for the mean. Both X alleles in females are represented in red and their sRNA mapping was divided by two for stochiometric comparison to the X allele in males (represented in green) and the Y allele in males (represented in blue). (I) Plot of X/Y genes DNA methylation in CHH context. Both X alleles in the female are represented in red, the X allele in the male in green and the Y allele in the male in blue. All X/Y genes were combined to plot the average proportion of methylated reads at cytosine positions along sliding windows; the 95% confidence interval is represented as a small ribbon around the curve. Twenty windows of 200 bp were studied upstream of the transcription start site (TSS) and twenty windows of 200 bp were studied downstream of the transcription termination site (TTS). The gene body (from TSS to TTS) was divided into twenty windows of equal size.

**Fig. 3.**
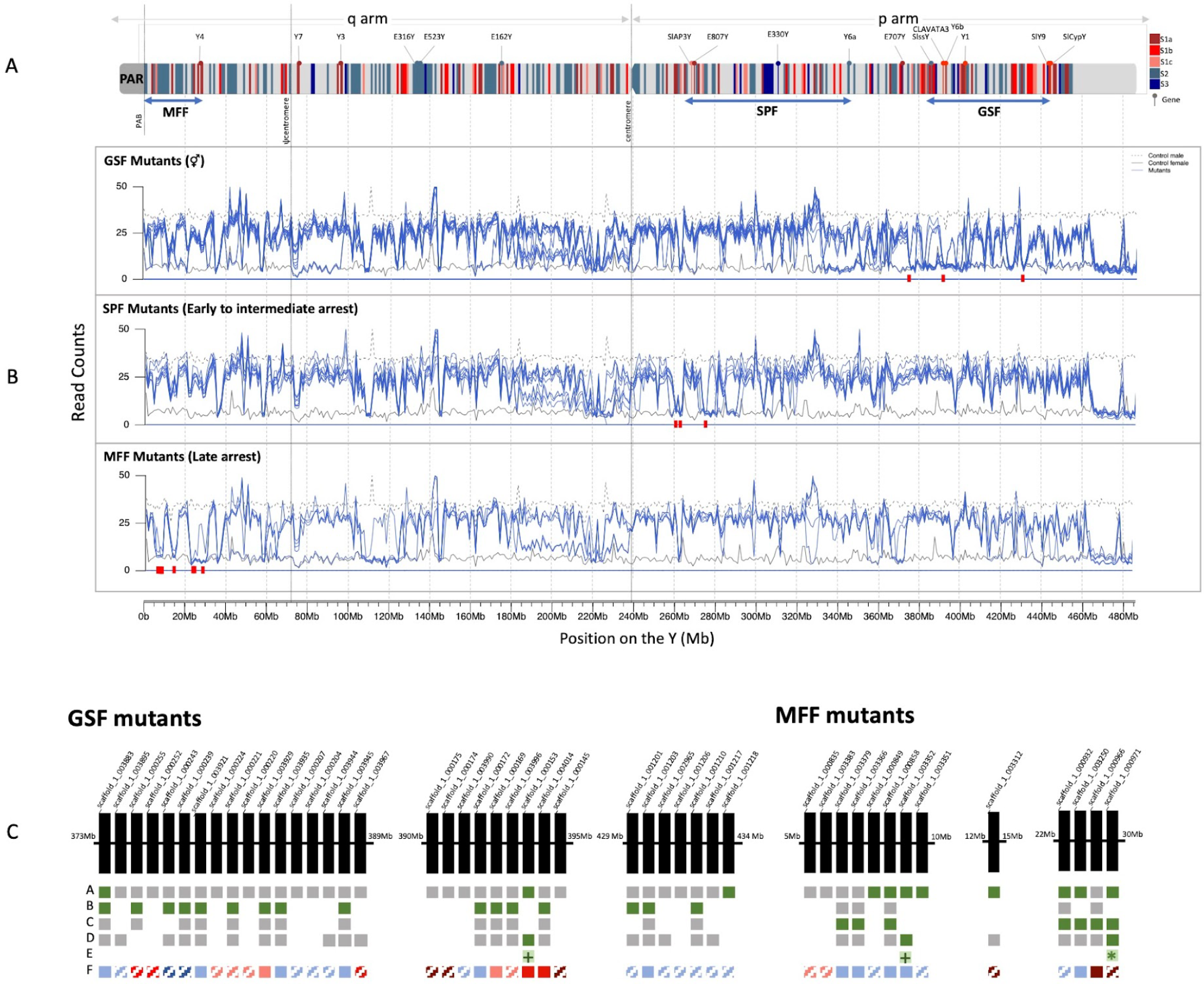
Y deletion mutants analysis. (A) The Y chromosome sequence and its putative sex-determining regions (MFF, SPF, and GSF) with estimated location from Bergero et al. (2008). The 417 gametologs genes are placed and colored according to their evolutionary stratum (S1a, S1b, S1c, S2 and S3). Additionally, genes previously characterized in the literature are identified by a ball-headed pin colored according to their stratum and located over the Y chromosome sequence. (B) Each plot shows the read count of all individuals (blue) grouped by mutant’s phenotypic category (from top to bottom: GSF mutants, SPF mutants, and MFF mutants) after mapping their reads onto the reference Y chromosome. The read count of the female and male control are also present in the plot (solid and dashed black line, respectively) and coincide with the maximum and the minimum (male and female values, respectively) read count of the mutants. Red rectangles depict genes that are deleted in all mutants of a given phenotypic category and present in all the remaining mutants (i.e., phenotype-specific deleted genes): 3 in GSF mutants at 370–440Mb, 3 in SPF mutants at 260Mb–280Mb and 8 in MFF mutants at 5Mb– 30Mb. The maximum number of reads was set to 50. Window size=1Mb. (C) Phenotype-specific deleted genes and their neighbor genes with a relevant presence/absence pattern within the low covered regions in the GSF and MFF mutants’ phenotypic category: row A concerns the presence/absence of the gene among mutants, in which green is for phenotype-specific deleted genes and gray is for genes deleted in at least one mutant of the category of interest and present in all the remaining mutants; row B and C concern the expression of each gene in a normal male at stage 5 and 8 of development, respectively: green is for genes expressed in the stage expected for sex determining genes (stage 5 for GSF genes or stage 8 for MFF genes), gray is for genes that do not follow the expected pattern of expression in sex determining genes, and blank means the gene is not expressed in any of the stages; row D concerns the functional annotation of each gene: green is for genes annotated with a sex determining function, gray is for genes annotated with no clear sex determining function, and blank is for genes without available functional annotation; row E regards they possible role as a sex determining gene, in which we highlight with asterisk (^*^) a very good candidate to sex-determining gene and with a plus signal (+) good candidates to sex-determining genes; row F indicates the stratum of each gene: brown for S1a, red for S1b, pink for S1c, light blue for S3 (for the genes with striped squares the stratum was inferred from the closest genes).

### Evolution of the sex chromosomes

The high-quality assembly of the X and Y chromosomes illuminates the evolution of recombination suppression between the sex chromosomes of *S. latifolia*. We found three evolutionary strata, and extensive rearrangements on the X, and especially the Y chromosome. Our analyses of rearrangements and estimates of synonymous site divergence (d_S_) between the X and Y copies used a set of 355 gametologs (= X/Y gene pairs) with 1:1 orthology with the outgroup species *Silene conica* and *Silene vulgaris*. Change-point analysis identified four regions in the non-recombining region with different pairwise mean X-Y d_S_ values in adjacent X chromosome regions, based on the gene order in the assembly (Figures 2 and S4A). We defined three evolutionary strata, S1, S2 and S3, based on the different d_S_ levels (Figures 2A and S4B). The oldest stratum, region S1, is split in two by S3. S3 has a lower d_S_ mean than its two flanking regions, which does not fit the expected pattern of decreasing divergence with proximity to the PAR; it has d_S_ value similar to that of the S2 region but much higher synteny, and it was not formed via the same type of rearrangements (see below). S1 and S3 are small regions located within the first 27 Mb of the X chromosome q arm. S2, on the other hand, is very large and includes most of the X chromosome. Using a molecular clock approach, we found that strata S2 and S3 evolved 4.9 [3.8, 6] and 4.4 [3.3, 5.6] MY ago respectively and stratum S1 10.8 [9.7, 11.9] MY ago, i.e. when the *S. latifolia* sex chromosomes originated (*18*).

Comparisons of gene order between *S. latifolia* and two non-dioecious close relatives, *S. conica* and *S. vulgaris* revealed large syntenic blocks with some rearrangements (Figure S5). The *S. latifolia* Y chromosome stands out as highly rearranged compared with either the X or both outgroups, with the notable exception of the S3 stratum, which includes in particular a syntenic block of 4 Mb (Figure S6A and S6C). The *S. latifolia* X shows homology with four

*S. vulgaris* scaffolds (1, 3, 6 and 16; Figures S5) and with chromosome 5 of *S. conica* (its closest relative), and smaller parts of chromosomes 1, 2 and 6 (Figures S5 and S6B). Reconstruction of the rearrangements between the X, the Y and the outgroups (Figures 2B, S6C-E and S7) showed that stratum S1 may have resulted from two inversions, one on the X encompassing S1a to S1b and one on the Y including S1c, that occurred early in the evolution of the sex chromosomes.

Stratum S3 had a lower d_S_ than stratum S1 and was the only region showing extended synteny between the X and Y chromosomes, which suggests that S3 has recently translocated into the middle of the oldest stratum S1 (Figure S6C). However, comparisons between *S. latifolia* X and the outgroups suggest that S3 was ancestrally located within the S1 stratum at the very same place (Figures 2 and S6D-E). To reconcile these findings, we propose that S3 (initially within the S1 stratum) was lost from the Y and later regained by a recent duplicative-translocation from the X (Figure 2B). Stratum S2 is likely slightly older than S3 as it is more rearranged, and probably arose through a different mechanism (Figure 2C). Reconstruction of the rearrangements between the X, the Y and the outgroups (Figures 2 and S8) could not associate S2 formation to a single event. We found several inversions, some of them pericentric, as previously speculated (*19*).

The Y and the X chromosomes are giant sized in *S. latifolia* and we infer that this is due to TE accumulation in both sex chromosomes. Our repeat analysis indeed revealed massive TE accumulation on the Y, but also, to a lesser extent, on the X (Tables 1 and S3, Figure 1). This explains why the X and Y, are, respectively, 4 and 5.5 times larger than a typical *S. conica* chromosome such as chromosome 5, with which most of the orthologs with the sex-linked genes are from. The *S. latifolia* autosomes average size is twice that of their *S. conica* homologs, supporting the view that TEs on sex chromosomes constitute a reservoir spreading genome-wide (*20*). In Eukaryotes, the non-recombining pericentromeric regions are TE-rich as recombination helps purge deleterious TE insertions (*21*) and this pericentromeric effect on the X chromosome is strikingly large (Figure S2B). The Y chromosome exhibits signs of considerable degeneration. Out of 1,541 1:1 orthologs in *S. latifolia* X, *S. conica* and *S. vulgaris*, 963 had no detected ortholog on the Y chromosome. A model-based phylogenetic analysis of gene gain and loss confirmed this observation. As many as 58% of the genes on the Y appeared to have been lost since it stopped recombining with the X about ∼11 MY ago. In addition, we detected more genes with premature stop codons on the Y compared to the X, except for stratum 3 (Figure 2E), suggesting the X might also be degenerating in this stratum. Among the gametologs with apparently functional X and Y copies, 77% of those with significant differences in rates of non-synonymous versus synonymous (d_N_/d_S_) changes between X and Y had the Y copies with higher d_N_/d_S_ values, indicating relaxed selection (Figure 2D). Another form of degeneration is when Y-linked genes have lower expression than their X counterparts, which has been reported in *S. latifolia* (*22-24*); this may be explained by epigenetic modifications (Figure 2F-I), as Y genes had more TEs in their vicinity and bear hallmarks of silencing, i.e., a higher number of 24 nucleotide small RNA mapping and higher levels of DNA methylation, especially around the promoter in the CHH context, compared to X genes (other contexts are shown in Figure S9). Using RAD-seq data, we found that the Y chromosome exhibits considerably lower genetic diversity *as* c*o*mpared to the X chromosome and autosomes (Figure S10), in agreement with this chromosome undergoing genetic degeneration .

### The sex-determining genes on the Y chromosome

We identified candidate sex-determining genes by sequencing at low coverage 18 mutants for sexual phenotypes (Table S4 and Material and Methods). Hermaphrodite mutants have deletions in the GSF region, asexual mutants in which anther development was stopped at early or intermediate stages in flower development have a deletions in the SPF region, and asexual mutants with pollen defects (late events) have deletions in the MFF region. SPF and MFF mutants were phenotypically very diverse, suggesting they have several different deleted genes (*25,26*).

Figure 3A shows the mapping of the mutant reads onto the reference Y chromosome along with reads from normal (U17) males and females, which indicate the expected coverage for presence (male) or absence (female) along the Y chromosome. Note that female coverage is generally low but not always null depending on how many X reads can mismap on the Y chromosome. A region is inferred as deleted in a mutant if the coverage is similar (or lower) to the normal female and different from the normal male. While the data are noisy, regions of very low coverage in all mutants of a given category (but not other categories) are still easily identified and are limited in number (3 for GSF; 2 for SPF; 3 for MFF categories). They were located as expected on the Y based on a map of the Y built using genetic markers to genotype the mutants and locate their deletions (*26*). In particular, MFF deletions cluster near the PAR and Y4; GSF deletions are close to Slss, DD44, Cyp and Y6a; SPF deletions are close to Y6b.

One of the three GSF deletions includes *Clavata3* (*scaffold_1_000153*), the recently proposed GSF candidate gene (*16*). Differences in the balance of the Clavata-Wuschel pathway in males and females has been proposed to explain carpel formation/inhibition in female and male flowers as *Clavata3* (a carpel inhibitor) has a functional copy on the Y and a pseudogene on the X, while *Wuschel* (a carpel promoter) is present on the X and deleted from the Y (*27*). Consistent with this, both *Clavata3* and *Wuschel* are in stratum 1, as expected if both changed during the first step of sex-chromosome evolution, i.e., they support the model involving male-sterility and female-suppressing mutations (*28*). Figure 3B focuses on genes located in the deletions, combining information on coverage, gene expression during early or late flower development (Table S5) and functional annotation of sterility terms. We also considered genes that are absent in at least one mutant of a category, possibly explaining the observed phenotypic variability observed among the mutants. We found several MFF candidate genes. A notable MFF candidate is the gene *scaffold_1_000971* that encodes a cytochrome P450 protein and is homologous to the *Arabidopsis thaliana* Cyp704B1 gene, which is crucial for pollen maturation, is expressed in the tapetum, and involved in the sporopollenin synthesis (a pollen cell wall component). Its inactivation causes male-sterility in *A. thaliana*. This gene is expressed in stage 8 but not stage 5 in *S. latifolia* male flower development consistent with a MFF gene. Another MFF candidate gene is *scaffold_1_003352* that encodes for a papain-like cysteine protease also expressed in tapetum and important for pollen maturation, through involvement in proteolysis and tapetal cell degeneration). It is also annotated as a male-sterility gene. No clear SPF candidate was found and this region is not shown in Figure 3B. *scaffold_1_000971* is probably located in stratum 1 while *scaffold_1_003352* is located in stratum 2. Our best MFF candidate is thus located in S1 as the best GSF one, which suggests that S1 might have formed when successive closely linked female-suppressing/male-enahncing mutations appeared during the evolution of dioecy (*28*).

## Conclusions

We produced high-quality *S. latifolia* sex chromosome assemblies, which provided insights about their structure, function, and evolution. We found that the evolution of the *S. latifolia* sex chromosomes started ∼11 MY ago with the differentiation of a small region including the GSF and MFF sex-determining genes. This first stratum probably formed by two paracentric inversions on both the X and the Y. More recently, another stratum including the centromere, S2, was formed ∼5 MY ago. This generated a very large non-recombining region on the Y, in which a massive accumulation of TEs occurred, and from which they dispersed throughout the chromosome. These repeats and the lack of recombination probably allowed chromosomal rearrangements to occur on the Y. The absence of recombination also led to genetic degeneration, with as much as 58% of the genes on the X being lost from the Y. Interestingly, similar changes also affected the X chromosome, as its non-recombining pericentromeric region also expanded to a giant size via massive TE accumulation, which might have been driven by a reduced effective population size of the X chromosome (*29,30*).

## Acknowledgments

We thank Elise Lacroix technician at Univ. Lyon 1 glasshouse for her help for growing the *S. latifolia* U17 plants.

## Author contributions

Conceptualization: GABM, RH, AMa, ADG, ABe, TG, DC, AMu

Formal analysis: CM, CB, QR, PV, PJ, AMu, MH, ET, XL, EMG, CL, PDF, CBe, RCe, RRV, GA, AEF, VM, AV, SK, ADG, AMa

Methodology: CM, CB, RL, ADG, GABM

Investigation: CM, CB, QR, PV, PJ, AMu, MH, ET, XL, EMG, CL, PDF, CC, KL, CBe, JB, SS, RCe, RL, RRV, GA, ABo, RT, DBS, SK, LFD, DC, TG, ABe, ADG, AMa, RH, GABM

Resources: RH, ADG, LFD, DBS, MSO, DRT, OAJA, MK, HS, HSi, PC, ZT, HT, LW, SD, BS, DP AMu

Software: CM, CB, ET, RL, RRV, AV, ADG

Validation: AC, RCo, CF

Visualization: CM, CB, QR, PV, AMu, ET, RL, RRV, DP, SK, AMa,

Funding acquisition: GABM, ABe Project administration: GABM

Supervision: GABM, RH, AMa, ADG, ABe, TG, LFD, SK, DBS, RCo, RT, ABo, RRV, RL, AMu

Writing – original draft: GABM, AMa, ABe, SK, DBS, SS, JB, PDF, XL, ET, AMa, QR, CB, CM

Writing – review & editing: GABM, TG, DC, LFD, SK, ABo, RRV, AMu, PV

### Data and materials availability

All data will be available on NCBI and codes on github.

## Supplementary Materials

Materials and Methods

Figs. S1 to S11

Tables S1 to S5

References (*31*–*102*)

