## Supplementary Material for "The *Silene latifolia* genome and its giant Y chromosome"

**The PDF file includes:**

Materials and Methods
Figs. S1 to S11
Tables S1 to S5
References 31-102

### Materials and Methods

Plant material

The *S. latifolia* U17 population (original U9 population gift of Sarah Grant, University of North Carolina, USA, described in (*31*)) was made by 17 generations of full-sib mating and used as an inbred line for all the experiments described. The U17 plants were grown at either LBBE, CNRS / Univ. Lyon 1 campus de la Doua, Lyon (for the genomic analysis) or IPS2, Univ. Paris-Saclay (for the transcriptomic analysis) in a glasshouse under controlled conditions optimal for *S. latifolia* growth following (*32*). Leaves (cut in pieces, genomic analysis) or flower buds (transcriptomic analysis) samples were collected and flash frozen in liquid nitrogen.

DNA extraction and sequencing

**DNA extraction:** 1g of frozen -80°C preserved leaves was ground in a mortar to a fine powder and transferred in a 50 ml Falcon tube containing 20 ml of CTAB Carlson buffer (100 mM Tris-HCl, pH 9.5, 2% CTAB, 1.4 M NaCl, 1% PEG 8000, 20 mM EDTA) including 10 mM DTT and 50 µl of RNase A (100 mg/ml solution). Following incubation at 65°C for 1 hour with occasional inversions, the sample was centrifuged to remove plant debris, chloroform/IAA extracted and gently mixed using a Hulamixer (Invitrogen). DNA was finally precipitated with 0.7 volumes of isopropanol.

**Purification and quality control:** The crude DNA extract precipitate was directly dissolved in G2 buffer and purified with QIAGEN Genomic-tip 100/G gravity columns (Blood and Cell Culture DNA Kit). The eluate was isopropanol precipitated and redissolved in LoTE buffer (10 mM Tris, 1 mM EDTA). Crude DNA extract precipitate was directly dissolved in G2 buffer and purified with the QIAGEN Genomic-tip 100/G gravity columns (Blood and Cell Culture DNA Kit). The eluate was isopropanol precipitate and redissolved in LoTE buffer (10 mM Tris, 1 mM EDTA).

**ONT library preparation and sequencing**: Small fragments were first eliminated from the genomic DNA by precipitation using the Short Reads Eliminator (SRE) kit or the Short Read Eliminator XL (SRE-XL) kit (PacBio, Menlo Park, CA 94025 USA). Libraries (two for each flow cell) were then prepared following the Oxford Nanopore « Genomic DNA by Ligation (SQK-LSK109) » protocol, DNA fragments (1 to 2µg) were repaired and 3'-adenylated with the NEBNext FFPE DNA Repair Mix and the NEBNext® Ultra™ II End Repair/dA-Tailing Module (New England Biolabs, Ipswich, MA, USA). Sequencing adapters provided by Oxford Nanopore Technologies (Oxford Nanopore Technologies Ltd, Oxford, UK) were then ligated using the NEBNext Quick Ligation Module (NEB). Aliquots of the libraries were mixed with the Sequencing Buffer (ONT) and the Loading Beads (ONT) then loaded onto R9.4.1 PromethION flow cells. During the run, each of the five flow cells was washed with DNase I once or twice following the « Flow Cell Wash Kit (EXP-WSH003) » and further aliquots of the libraries were loaded. Two additional libraries were prepared in the same way, but using the SQK-SKQ110 kit (following the Oxford Nanopore « Genomic DNA by Ligation (SQK-LSK110) » protocol) and one aliquot was loaded onto one R9.4.1 PromethION flow cell. The other aliquots of the libraries were used for reloading the flow cell after two DNase I treatments during the run. Reads were base-called using Guppy version 3.2.10+aabd4ec for the five first runs and Guppy version 4.0.11+f1071ce for the last one.

**Illumina library preparation and sequencing**: A PCR-free library was prepared using the Kapa Hyper Prep Kit (Roche, Basel, Switzerland). Genomic DNA (1.2µg) was sonicated using a

Covaris E220 sonicator (Covaris, Woburn, MA, USA). Fragments were end-repaired and 3′-adenylated, and Illumina adapters (Bioo Scientific, Austin, TX, USA) were ligated according to the manufacturer's instructions. Ligation products were purified twice with AMPure XP beads (Beckman Coulter Genomics, Danvers, MA, USA) and quantified by qPCR (MxPro, Agilent Technologies, Santa Clara, CA, USA) using the KAPA Library Quantification Kit for Illumina Libraries (Roche). Library profile was assessed using an Agilent High Sensitivity DNA kit on the Agilent 2100 Bioanalyzer. Library was paired-end sequenced on an Illumina NovaSeq 6000 instrument (Illumina, San Diego, CA, USA) using 151 base-length read chemistry.

Short Illumina reads were bioinformatically post-processed following (*33*) to filter out low quality data. First, low-quality nucleotides ($Q < 20$) were discarded from both read ends. Then remaining Illumina sequencing adapters and primer sequences were removed and only reads ≥ 30 nucleotides were retained. These filtering steps were done using in-house-designed software based on the FastX package (*34*). Finally, read pairs mapping to the phage phiX genome were identified and discarded using SOAP aligner with default parameters (*35*) and the Enterobacteria phage PhiX174 reference sequence (GenBank: NC_001422.1).

**Hi-C library preparation and sequencing**: The Hi-C library was prepared using the Dovetail Omni-C kit (Dovetail Genomics, Scotts Valley, CA, USA), following the commercial protocol for non-mammalian samples version 1.0. Briefly, flash-frozen young leaves (300 mg) were cryoground in liquid nitrogen; the chromatin was fixed with formaldehyde (only), randomly digested with DNase I and then extracted. Chromatin ends were repaired and ligated to a biotinylated bridge adapter, followed by proximity-ligation of adapter-containing ends. After proximity ligation, crosslinks were reversed and DNA was purified. Purified DNA was treated to remove biotin that was not internal to ligated fragments, and a sequencing library was generated using NEBNext Ultra enzymes and Illumina-compatible adapters. Biotin-containing fragments were isolated using streptavidin beads before PCR enrichment of the library. The Dovetail Omni-C library was sequenced on an Illumina NovaSeq6000 instrument (Illumina, San Diego, CA, USA) using 150 base-length read chemistry in paired-end mode, producing more than 200 M reads.

**Optical map**: High Molecular Weight (HMW) DNA was prepared from 2 g of flash-frozen young leaves, according to the following protocol https://www.protocols.io/view/isolation-and-extraction-of-plant-nuclei-in-plug-6qpvr6632vmk/v1. The Direct label and stain (DLS) labeling (DLE-1) protocol was performed according to the Bionano Genomics (San Diego, CA, USA) instructions, starting with 750 ng HMW DNA. The labeling preparation was loaded onto a chip as recommended by Bionano Genomics.

Genetic mapping

**Plants for mapping crosses**: Seeds collected from plants growing in the wild near Zagreb, Croatia (in 2000) and near Xativa, Spain (in 2010) were grown in the greenhouse at Indiana University and within-population crosses were made to produce seeds for experimental studies. For this study, two females from Croatia were each crossed with a different male from Spain, to produce a maternal F1 and paternal F1 plant. The F2 offspring of these two independent interpopulation crosses were used for genetic mapping. The offspring were grown in greenhouse conditions at Indiana University and in field conditions in Spain and Croatia. The greenhouse plants were grown in a 50:50 Metromix (Scotts-Sierra Horticultural Products, Marysville, OH) greenhouse soil mixture, and transplanted to 12.7 cm pots after developing 4 leaves, and kept at ~24$^{o}$C in 16h:8h light:darkness cycle to induce flowering. DNA was collected by freezing portions of leaves and later grinding them in liquid N2 and processing the resulting powder with a Dneasy Plant mini kit (Qiagen) following the manufacturer instructions.

**SNP assay and genotyping:** A SNP assay was designed based on DNA sequencing of the parental plants of the cross using 150 bp paired-end reads on a NextSeq500 (Illumina inc) at the Centre for Genomics and Bioinformatics at IU. In total we generated 296,553,201 reads, which were trimmed for adapters and low-quality bases using Trimmomatic v0.32 (*36*).We called SNPs by mapping the paired sequenced reads to the reference *S. latifolia* genome from (*37*) using Bowtie2.2.9 (*38*); read pairs with a MAPQ of less than 10 were discarded. SNPs were identified using mpileup from Samtools v1.3.1 (*39*) and varscan v2.4.2 (*40*). These SNPs were filtered for coverage and designated as new or known based on their presence in scaffolds with known genetic map locations (*37*). An assay was designed using Design Studio Microarray Assay Designer (Illumina Inc) with 5,000 previously mapped and 5,000 novel SNPs. Genotyping was performed at the Van Andel Research Institute (Michigan) and Research Technology Support Facility Genomics Core at Michigan State University using an Illumina Infinium platform. We genotyped the parents and mapping population grown in both the greenhouse and the field using Illumina GenomeStudio V2011.1 (Illumina Inc). SNPs were retained if they showed clear clustering of two distinct homozygous genotypes and one group of heterozygous individuals and if their minimum call rate was 0.98.

**Genetic map construction**: The SNP calls from 183 females and 307 males from one family were exported and formatted for use with Lep-MAP 3 (*41*). Briefly we did not employ filtering as it is not recommended for mapping a single family, SeparateChromosomes2 was run with distortionLod=1 and lodLimit=10, and OrderMarkers2 was used to construct a sex-averaged, as well as separate male and female recombination maps. The expected number of 12 linkage groups was obtained. Comparison of the sex-specific maps allowed us to identify the pseudoautosomal region (PAR), which is linked to the fully X-linked region (here termed "X-NR") but showed recombination in both sexes. We compared our genetic map with that of (*37*) based on SNPs in scaffolds that were mapped in both studies, and made them as collinear as possible. 3 markers mapped to different chromosomes and were removed from the final map. The markers mapped by (*37*) are also largely collinear with those in the map published by (*42*).

<u>Assembly, scaffolding and anchoring to the genetic map</u>

**ONT contigs**: The assembly strategy was to first create a draft assembly using the long reads and the Flye assembler (*43*). This first assembly was improved by a polishing step using the short and long reads in order to correct errors and improve consensus quality (*44*). We separated polishing into three parts: one using the long reads with the software Racon (*45*); then using Illumina reads using the software ntEdit and nthit completing three rounds at k=40,50,60 (*46*); a final step used the short read alignments with the software pilon (*44*) (Figure S1). These steps were iterated several times to generate an accurate consensus sequence.

We generated an initial polished long-read assembly with a total size of 3.7 Gb and a N50 of 8.7Mb (Supplementary Table S1) and a maximum contig size of 87 Mb. BUSCO (*47*) metrics were calculated to evaluate the number of conserved genes using the Viridiplantae database (435 genes), and the gene completeness analysis shows that our polished assembly achieves a 99.1% (421/425) (Table S2). The resulting assembly is longer than the size described in the literature (2.6-3.2 Gb) (*9*). Visual inspection of the assembly graph suggested duplication of sequences due to the heterozygosity of parts of the genome, which was confirmed by the SNP density analysis later on (Figure 1A). We compared the assembly's contigs within them and removed the duplicates in the assembly using the software purge haplotigs (*48*). The reduced assembly size was 2.7 Gb, of which 1 Gb was associated with alternative haplotypes with an N50 of 16.5Mb, showing that the larger contigs were conserved after the genome reduction. Following contig assembly, we used Bionano data.

**Bionano scaffolding**: Following the contig assembly, we used Bionano data to generate scaffolds and error-correct contigs. The DLE-1 preparation was run on a single flow cell. The molecules generated were assembled to produce an optical map by software provided by Bionano Genomics (BNG) with the following two options: "add pre-assembly" and "non haplotype without extend and split" (bionano solve and tools Version: 3.6.1). The optical map was filtered to keep the maps corresponding to just one haplotype. The optical map and the ONT contigs were scaffolded by the hybrid scaffolding pipeline to generate the hybrid scaffolds. BisCoT was used to correct artifactual duplications (negative gaps) introduced during the scaffolding process (*49*). A final polishing step was performed using Hapo-G (*50*), using Illumina sequencing data. The assembly consisted of 1,274 hybrid scaffolds with a cumulative size of 2.8 Gb (Table 3).

**Hi-C scaffolding**: The bionano assembly was then integrated with the Omni-C data using the software SALSA (*51*) to further orient and order the scaffolds into super-scaffolds to reach a chromosome arm level assembly. We obtained 1,004 scaffolds with an N50 of 84Mb, reaching almost chromosome-arm length scaffolds.

**Anchoring to the genetic map**: To obtain a chromosome-level assembly, we used the separate male and female genetic map information (see above section "genetic map") to orient and order the scaffolds and reconstruct chromosomes using the software ALLMAPS (*52*). We first mapped all 10.000 markers to the assembly and selected the ones matching with good quality ($q > 20$; 7.916 markers) to ensure the correct positions. We anchored 12 chromosomes with a total size of 2.2 Gb, and then performed a manual curation process to correct inversions and translocations, and maximized the collinearity between the scaffolds and the genetic markers. There were a total of 710 unplaced scaffolds, totalling 576 Mb, which we used to build the Y chromosome assembly.

We used the information from Omni-C data to rearrange the scaffolds into the sex chromosomes (Figure S3).

Annotation

**Gene annotation**: We used two complementary methods to predict genes in the new reference genome: Braker v2.1.6.20 (*53*), which uses RNAseq data, and Helixer (*54*), which takes advantage of publicly available databases. Before gene prediction, we softmasked the genome with RepeatMasker (*55*) following de novo repeat detection with RepeatModeller (*56*). Prior to running Braker, we first trimmed short RNAseq reads derived from 12 samples and tissues (see section "Transcriptomic analysis" for details) with FASTP (*57*) to a length of 130, and then mapped them to the reference genome using gsnap (*58*) with standard parameters for RNAseq data. We merged each individual RNAseq alignment of each individual with Samtools (*39*), and then filtered the pooled dataset with the filterBam from Augustus (*59*) in order to keep unique mappings. The resulting alignments bam file was then used as input data and fed into Braker v.2.1.6 using Augustus for training and Diamond to filter redundant training gene structure (*60*). Second, Helixer (*54*), a machine learning program, was used on a GPU (Graphical Processing Units) with default parameters and using land plants database as learning examples.

We assessed the completeness of each of the two genome annotations independently with BUSCO v5.4.6 against the Viridiplantae database (*47*). Both annotations gave similar BUSCO scores, but an excessively high number of genes were predicted (80,140 by Braker and 52,672 for Helixer) including a high proportion of previously softmasked genomic regions. To reduce the rates of putatively false positive predictions, we used the intersection between the gene sets from the Braker and Helixer pipelines. The quality of this new prediction with BUSCO was as high as the two more exhaustive predictions. We finally manually added a few complete BUSCO genes present only in the Braker annotation.

**Functional annotation**: We recovered the protein fasta file from the combined annotation of Braker and Helixer, to obtain a functional annotation (Gene Ontology), with PANNZER2 (http://ekhidna2.biocenter.helsinki.fi/sanspanz/), using the default parameters (*61*). In total, 35,459 protein sequences were predicted and 11,816 of them were annotated by PANNZER2 with at least one function.

**Repeat annotation**: The EDTA pipeline (*62*) was run to find LTR retrotransposon superfamilies and to annotate DNA TEs, MITEs and Helitrons. LTR retrotransposons were further fine-annotated to family level using DANTE (https://github.com/kavonrtep/dante/). In total, we detected 40,824 full-length retroelements with at least one protein domain annotated. We complemented our repeatome study using the RepeatMasker v. 4.1.0. (*55*) tool to detect (i) simple and low complexity repeats and (ii) satellite, LINE repeats and 5S and 45S rDNA using a previously obtained *S. latifolia* specific library (*63,64*).

Genomes of *Silene conica* and *Silene vulgaris*

The sequencing, assembly, and annotation of the *S. conica* genome was recently described, based on a combination of PacBio HiFi sequencing, Hi-C sequencing, and Bionano optical mapping (*65*). The annotation pipeline further incorporated expression data from PacBio Iso-Seq and Illumina RNA-seq. The *S. vulgaris* genome was sequenced and assembled using the same combination of PacBio HiFi, Hi-C, and Bionano and assembled/annotated with the same bioinformatic pipeline as the *S. conica* genome. The *S. vulgaris* data were generated from a genotype that resulted from crossing genotypes sampled from Krasnoyarsk, Russia (KrIIId-11) and Kováry Meadows, Czech Republic (KOV52-4) and subsequent selfing of the F1 for a single generation. *Silene conica* and *S. vulgaris* data will be made available under the NCBI BioProject IDs PRJNA904366 and PRJNA1012482, respectively.

Analysis of evolutionary strata

We used orthofinder (*66*) to identify orthologous groups in the genomes of *S. latifolia*, *S. conica* and *S. vulgaris*. We also identified *S. latifolia* gametologs (i.e. alleles of orthologous genes present in the X and Y sex chromosomes) i using reciprocal best-blast-hits with Blastp (*67*) with an initial e-value of 1e-06 and the results from orthology reconstruction, using a set of 366 single copy orthologs in *S. conica, S. vulgaris*, and the X and Y chromosomes of *S. latifolia*. We computed synonymous divergence ($d_S$) per synonymous site and non-synonymous ($d_N$) divergence between gametologs and their standard errors (SE) using yn00 from PAML (*68*) on codon-based alignments produced by TranslatorX (*69*), using MUSCLE (*70*) as the protein aligner. In order to objectively identify the limits of evolutionary strata, we looked for changes in mean $d_S$ values along the X chromosome gene order (the most likely ancestral gene order) with a changepoint analysis performed using the R package mcp (*71*). This analysis relies on Bayesian regressions to infer the location of changes in means of the $d_S$ values. The MCMC was run for 100,000 iterations and used 3 replicates. We statistically compared a null model of a simple linear regression to a model with a single changepoint (i.e., two strata) against a model with two or three changepoints (three or four strata) using the loo R package. We also tested the significance of the differences in mean $d_S$ values, between pairs of adjacent strata, again using the X gene order, using the Savage-Dickey density ratio to compute the Bayes Factor (BF) and posterior probability of the difference. Specifically, we tested the two following hypotheses: i) the $d_S$ value of stratum 1 is greater than the value of stratum 2; ii) the dS value of stratum 2 exceeds that of stratum 3. We also tested this hypothesis using a TukeyHSD test based on an ANOVA as implemented in R.

As rough estimates of the age of evolutionary strata, we used the formula Tgenerations = $d_S/2\mu$, where $\mu$ is the mutation rate per bp and per generation, estimated to be $7.31 \times 10^{-9}$ mutations/bp/generation in *S. latifolia* (*18*) and an average generation time of 1.5 years (*72*). We used the mean, median, as well as the 15% highest quantiles of $d_S$ values observed between X and Y genes in each of the identified strata.

<u>Phylogenetic dN/dS analysis</u>

The previous analysis identified 366 orthologous genes in the X and Y chromosomes of *S. latifolia* (gametologs), *S. conica* and *S. vulgaris* (which were used as outgroups) with OrthoFinder (*66*) and its tools. We used MACSE v2 (*73*), to align the coding sequences (CDSs) of these genes based on their amino acid translation, taking account of the reading frame. Then, we ran PAML (*68*) with an unrooted phylogenetic tree in which the X and Y chromosomes of *S. latifolia* were treated as two separate lineages. We conducted analyses with branch and branch-site models. For analyses under branch models, we tested two alternative models: (i) a single value of $d_N/d_S$ ($\omega$) for the X and Y chromosomes genes, and (ii) different values of $\omega$ for the X and Y lineage. In all cases, *S. conica* and *S. vulgaris* were background branches (*74*). We then performed a likelihood ratio test (LRT) to compare the models. For the analyses under branch-site models, we also tested two alternative models: (i) the Y chromosome lineage was the foreground and the X chromosome, *S. conica* and *S. vulgaris* lineages the background, and (ii) $\omega$ was fixed at the value of 1 for the Y chromosome for sites under positive selection (null hypothesis). These branch-site analyses allow us to estimate the proportions of sites in each lineage that are under neutral, purifying, and diversifying selection (see PAML manual for details, see (*68*)). Note that the second approach removes false positive sites that are nearly neutral or under weak constraints in the foreground lineages (*75*). Then we collected the log-likelihood scores under both models to conduct the LRT.

<u>Quantification of gene gain and loss in a phylogenetic context</u>

We first explored the amount of gene gains and losses using a simple approach based on orthology inference, in which we investigated, for 1:1:1 orthologs among *S. vulgaris: S. conica: S. latifolia X,* how many genes had 0,1 or more orthologs on the Y, and counted the resulting gene gains and losses on the Y.

To further quantify gene gains and losses between the X and Y statistically in a phylogenetic context, we estimated a birth-death parameter ($\boldsymbol{\lambda}$) using the software Cafe version 5 (*76*). This requires a matrix of gene counts per species per gene family, and an ultrametric tree. We used the gene families inferred by OrthoFinder and reconstructed a species tree using all single-copy orthologs shared by *S. vulgaris, S. conica, S. latifolia X*, and *S. latifolia Y.* We aligned all the single-copy ortholog sets using MUSCLE (*70*), and constructed a supermatrix of all the alignments. We then estimated a maximum likelihood tree for the concatenated set of aligned genes using RAXML-NG (*77*) with 10 randomised parsimony starting trees, a fixed empirical substitution matrix (LG), inferred empirical amino-acid frequencies from the alignment, and eight discrete GAMMA categories. We performed 400 bootstrap replicates and assessed bootstrap convergence using the bsconverge option and a 1% cutoff. We compared this model to a model using an analysis partitioned by gene, providing the same evolutionary model as above (*i.e.* fixed empirical substitution matrix (LG), empirical amino acid frequencies from the alignment, eight discrete GAMMA categories) but for each gene independently. Given that the AIC of this more complex model did not suggest better support than for the simplest, less parameterized model, we used the former model, assuming the same sequence evolution model for all genes. The resulting maximum likelihood tree was provided as input to the R8s program. The calibration was set to 8 million years, based on the median divergence time estimated between *S. vulgaris* and *S. conica* and inferred using TimeTree (*78*).

The cafe5 program was run after excluding gene families with more than 100 copies, as families with large variance may not be informative, following the author's recommendations (*76*). We estimated an error model from our empirical data, to take account of genome assembly errors, and used it as input. We ran a model assuming a Poisson distribution for the root frequency distribution (a uniform distribution was also tested and provided lower log likelihood values). We incorporated among-gene-family rate variation using the Gamma model with K = 2 categories after checking that a simpler model with K = 1 or a more complex model with K > 2 categories did not fit as well, or produced a similar log-likelihood. Ten replicate runs were performed for each tested model.

The results were summarized using the CafePlotter (https://github.com/moshi4/CafePlotter) to obtain the total number of gene family expansions (additions of genes to the X or Y) and contractions (gene losses from the X or Y) on each branch of the tree, as well as the numbers that were statistically significant.

Analysis of chromosomal rearrangements

**Synteny analysis**: We first aligned the X and Y chromosomes of *S. latifolia* using Minimap2 (*79*). The resulting paf files were visualized using pafR to examine global synteny. An ideogram was then constructed using the RIdeogram R package in order to test the extent of gene duplication of a set of 7 key Y genes identified in this study. The same procedure was repeated, but mapping the Y genes to the X chromosome of *S. conica* instead of *S. latifolia.*

We then performed a GeneSpace analysis (*80*), to look for synteny and infer gene copy number variation among the reference genomes of *S. latifolia, S. conica* and *S. vulgaris*. This approach relies on blast, OrthoFinder and MCScanX to infer synteny among the genomes. The results were visualized using riparian plots as implemented in the R package GeneSpace (*80*). To further quantify the extent of rearrangement between the Y and X the GeneSpace analysis was repeated using only the X and Y chromosomes of *S. latifolia* as inputs.

**Scenario for chromosomal rearrangements**: We selected all orthogroups with exactly one exemplar in *S. conica* chromosome 5, *S. vulgaris* scaffold 1, *S. latifolia* chromosome X and *S. latifolia* chromosome Y. To avoid (as far as possible) errors in the orthology assignment, or retrotranspositions of genes between the X and Y, we analysed only genes whose immediate neighbor (no intervening genes in the dataset) in *S. conica*, *S. vulgaris* or *S. latifoli*a Y, is also a neighbor in the *S. latifolia* X chromosome. In addition we removed all the genes identified as belonging to strata S2 because we suspect that this region has a different, recent history than the rest of the chromosome. This resulted in 79 orthogroups of 4 genes.

We used Badger (*81*) to reconstruct the two ancestral states, the ancestral chromosome before the speciation from the outgroups, and the ancestral chromosome before the divergence of the X and Y chromosomes. We generated 1000 ancestral arrangements to compute a posterior probability for each. To infer inversions, we used DCJ2HP (*82*). We ran once for each of the five branches of the phylogenetic tree being considered. We generated 1000 different scenarios to allow for a diversity of possible scenarios.

In addition we simulated 1000 inversion scenarios starting from a virtual ancestral chromosome, using the same breakpoints as the inferred X-Y scenarios, with as many inversions as the inferred X-Y scenarios, in order to compare the properties of the simulated and inferred scenarios.

<u>Genetic diversity and differentiation analysis</u>

To investigate the genomic landscapes of diversity and differentiation in *S. latifolia* and between *S. latifolia* and the closely related *S. dioica*, we selected, two *S. latifolia* populations from France - (FRA1, 9 females, 6 males), and from Poland (POL3, 9 females and 8 males) and one *S. dioica* population from Belgium (BEL1, 10 females and 9 males) from a previously published dataset (*83*). Individuals were sequenced using 125 bp paired reads by the double digest restriction-site associated DNA (ddRAD) approach (see details in (*83*)). Raw reads were demultiplexed using the 'process_radtags' function in Stacks2 (*84*), and low-quality bases and adaptor sequences were trimmed using FASTP (*57*) with the following settings: a base phred quality (Q) of 15 (-q), a maximum of 50 % of bases with Q < 15 (-u), a base correction with a minimum length overlap of 60 bp (-c), for detection and removal of adaptors, and a per read cutting sliding window by quality score with a window size of 5 bp (-W) and a mean quality threshold of 15 (-M).

For analysis of autosomes and the X chromosomes, we mapped reads from female samples to the *S. latifolia* reference assembly after excluding Y-linked scaffolds using BWA-MEM (*85*) and performed genotype calling in diploid mode based on reads with mapping quality scores of at least 20 and bases with quality scores of 30 using freebayes (*86*). To study the Y chromosome, male and female samples were mapped to a complete reference assembly, including Y-linked scaffolds, but masking the PAR. We excluded X-homologous loci on the Y chromosomes ; these were identified as regions to which female reads could be mapped (covered by at least three reads from at least two females). Genotype calling on Y-linked scaffolds was conducted in males as described above, but in haploid mode. In both males and females, genotype calls supported by less than six reads were masked, and sites with genotyping rates below 50% in each population or showing low and high allelic balance "AB < 0.25 or > 0.75" were discarded (*87*).

We subsequently used the python package pixy (*88*) to calculate the following population genetic statistics: weighted Hudson's $F_{ST}$and nucleotide diversity ($\pi$) based on RAD-tags that included at least 100 bps after the above filters. To investigate patterns in $F_{ST}$ and $\pi$ along autosomes and the X chromosome, we calculated means in 5 MB windows containing between 30 and 107 RADtags, after removing windows with fewer than 5 RADtags. For the autosomes and X chromosome, we used female samples only, whereas we used only males for the Y-chromosome.

<u>Epigenetic analysis</u>

**Small RNA sequencing:** small RNAs were extracted and purified using the Sigma mirPremier microRNA Isolation Kit (reference SNC10, small RNA protocol) from calyx samples dissected from 4mm-long flower buds and from young leaves from 3 females and 3 males of *Silene latifolia* grown in controlled greenhouse conditions in Lyon, France. The sequenced individuals were brothers and sisters of the individuals used for genome sequencing. Small RNA integrity was checked before sequencing with the fragment analyzer. sRNA libraries were then produced using the NEXTflex Small RNA-Seq Kit v3 from Bioo Scientific. Sequencing was carried out on an Illumina single-end NovaSeq 6000 platform by Montpellier Genomix (https://www.mgx.cnrs.fr).

**small RNA data analysis:** The sequencing company evaluated the quality of the raw sequencing data using FastQC (*89*) and removed adapters and trimmed low-quality reads using Cutadapt (*90*). Only reads of length 24 nt were used for subsequent analyses as they are known to be mostly involved in driving DNA methylation. ShortStack v4.0.0 (*91,92*) was used to map 24nt small RNA reads to the *S. latifolia* reference genome. Female samples were aligned to a genome without the Y chromosome to avoid aberrant mapping of female small RNA to the Y chromosome. The number of small RNAs mapping to each gene was extracted with BEDtools (*93*) from the transcription start sites (TSS) to the transcription termination sites (TTS) plus 200 bp upstream of the TSS for the promoter.

**Bisulfite-sequencing (BS-seq):** one male and one female full sibs of the individual whose genome was sequenced) were sampled for flower bud characterization of DNA methylation through BS-seq. 4mm buds were dissected for the calyx and snap frozen in liquid nitrogen. DNA was extracted using the Qiagen Kit DNeasy Plant MiniKit. The DNA was treated with RNAse and quantity and quality were checked with the Fragment Analyser. Libraries were prepared using the Swift Bioscience ACCEL-NGS Methyl-SEQ DNA library kit. 0.1% of unmethylated phage Lambda DNA was added. The DNA was then sonicated to obtain an average fragment size of 350 bp. The DNA was bisulfite converted using the Zymo Research EZ DNA Methylation Lightning kit. A low-complexity (GC rich) 10 nt polynucleotidic tail was added with an adaptase and primers were ligated, followed by six PCR cycles. Sequencing was then performed by Montpellier Genomix on an Illumina Nova-Seq paired-end S1 flowcell with 150 nt long reads.

**BS-seq data analysis:** The sequencing company used the Phage lambda DNA to estimate the non-conversion rate of the experiment using Bismark (version 0.22.1). BS-seq reads were cleaned and trimmed with trim_galore (version 0.6.8dev, options --paired --clip_R1 2 --clip_R2 2 --three_prime_clip_R1 2 --three_prime_clip_R2 2 --illumina, see (*94*)) and quality was checked with FastQC v0.11.9 (*89*). Bismark version 0.24.1 (*95*) was then used to map reads to the genome with Bowtie2 version 2.5.1 (*38*). Only unambiguously mapped reads were retained. The resulting bam files were sorted with Samtools sort version 1.10 (*39*), mates were then fixed with Samtools fixmate, the files were sorted again, then duplicates were removed with Samtools markdup (option -r) and the files were sorted again. Finally, methylation calls were extracted with Bismark methylation extractor (options --CX_context --no_overlap). Gene metaplots were drawn by combining every gene's raw proportion of methylated reads at cytosine positions from 4000 bp upstream of the gene to 4000 bp downstream. The region was divided into 20 windows of 200 bp upstream and downstream of the gene, and the regions between the genes' TSS and TTS were divided into 20 windows of equal size. For the circosplot, the genome was cut into windows of 1 Mb and the raw proportion of methylated reads at cytosine positions was computed in each context and for the male and female individual.

**Gene expression analysis:** public RNA-seq reads from *S. latifolia* for four males and four females that were sampled for leaves and flower buds (see (*32*); ENA project PRJEB14171) were used for gene expression estimates. Reads were mapped to annotated transcripts with the software Kallisto (*96*). Female samples were mapped to transcripts after excluding Y genes. X-Y gene pairs identified in this study were used to estimate the ratio of Y to X gene expression in males.

Analysis of mutants with Y deletions

We re-sequenced 17 individuals previously characterized (*25,26*) that carry deletions of Y chromosome regions that affect the sex phenotype (resulting, for example, asexual or hermaphrodite individuals) (Table S4). The genomic DNA of one mutant (MH115) was amplified to increase the amount of DNA available for sequencing using the GenomiPhi V2 DNA Amplification Kit from illustra.

A PCR-free library was prepared using theTruSeq DNA PCR-Free Kit (Illumina, San Diego, CA, USA). Genomic DNA (0,5 to 1µg) was sonicated using a Covaris E220 sonicator (Covaris, Woburn, MA, USA). Fragments were end-repaired and 3′-adenylated, and Illumina indexed adapters were ligated according to the manufacturer's instructions. Ligation products were purified twice with AMPure XP beads (Beckman Coulter Genomics, Danvers, MA, USA) and quantified by qPCR on a LightCycler 96 Real-Time PCR System (Roche) using the KAPA Library Quantification Kit for Illumina Libraries (Roche). Genomic library was controlled (sizing and estimation of the concentration) by electrophoresis on a AATI Fragment Analyzer™ (Advanced Analytical Technologies, Ankeny, IA, USA) device with the DNF-474 High Sensitivity Fragment Analysis Kit**.**

We sequenced the individuals on the Illumina NovaSeq 6000 platform to obtain 150 bp paired-end reads. replicates were obtained for 7 individuals. Thus our final dataset comprises 24 samples. As a control, we added a male and a female individual with normal phenotypes (the same individuals whose short reads were used to generate the reference male genome). All the samples were filtered with FASTP using the default parameters 6 (*57*).

We mapped the reads from the mutants and controls to the masked male reference genome using BWA-MEM (*85*), which was previously shown to be efficient for mapping plant genomes (*97*). Data processing was performed with SAMtools ver1.16.1 (*39*), including filtering by mapping quality >=30. To identify deleted genes in the mutants and select candidates for sex-determining genes we calculated the mean depth per base for each gene in the gene annotation step per mutant or control individual. We considered the mean depth of the female control as the minimum value for a gene to be considered present. Further, to consider only Y-linked genes we filtered out genes that were covered by reads from the control female kept genes that were present in our control male. We grouped the absent genes according to the mutants' phenotypic category (hermaphrodite or asexual with arrest of anther development at different stages). For our initial list of candidate genes that might affect each phenotype, we selected genes that are absent in all the mutants of a given category and are present in all the remaining mutants. Finally, we looked at the neighboring genes of our initial list of candidate genes that are missing in at least one mutant of a given phenotypic category and are present in all the remaining mutants to consider the possibility that more than one gene plays a role in the phenotypic categories.

Transcriptomic analysis of floral development

Flower staging was carried out as described in (98). We focused our transcriptome analysis on flowers at stage 5 and stage 8. At stage 5, all of the flower primordia are initiated. At stage 8, male and female flowers can be easily distinguished. Three biological replicates were analysed for each stage. Total RNA was extracted from frozen flowers using RNeasy plant mini kit (qiagen ref:74904). RNA quality and concentration were evaluated with a Bioanalyser 2100 (Agilent Technologies) on Agilent RNA Pico chips. RNA recovery ranged from 900–2000 pg/µl, with a RNA Integrity Number (RIN) above 7. RNA sequencing (RNA-seq) libraries were constructed according to Illumina instructions and sequenced on a HiSeq 4000 platform at Novogene Co. Ltd. Raw sequencing reads were cleaned by removing adaptor sequences and low-quality reads (Table S3). The resulting high-quality reads were mapped to the S. latifolia genome, using STAR version 2.7.5c (*99*), with the default parameters. RNA-seq from the male flowers were mapped to the *S. latifolia* genome assembly including both the X and Y chromosomes. RNA-seq from the female flowers were mapped to the *S. latifolia* genome assembly excluding the Y chromosome. Gene expression was quantified using featureCounts with default parameters (*100*). Read count normalization and differential expression analysis were performed using DESeq2 (*101*). Differentially expressed genes (DEGs) selected with adjusted P-value $\leq 0.05$ and log2 FC $< -0.5$ and $>0.5$ were filtered for subsequent analyses. Analysis of GO term enrichment was performed using ClusterProfiler (*102*).

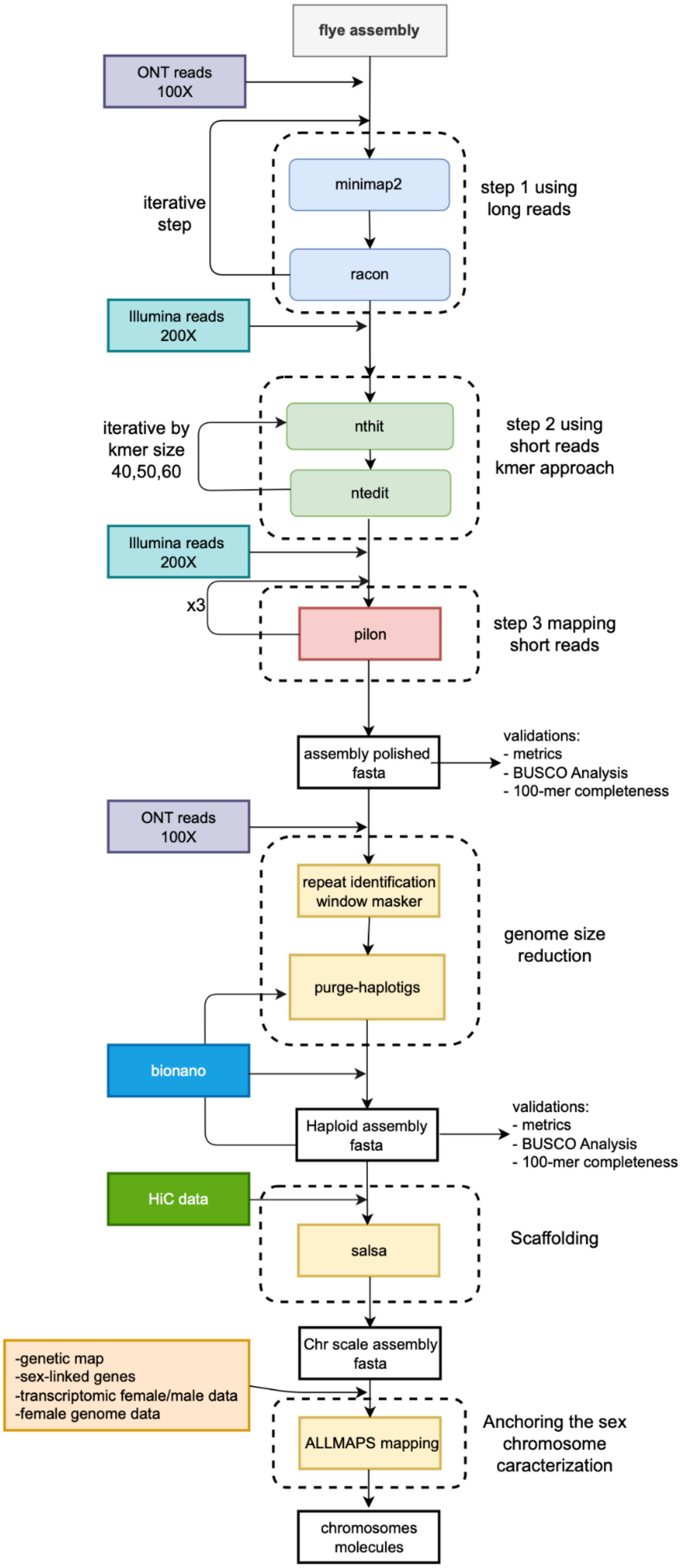
flye assembly
ONT reads
100X
iterative
step
minimap2
racon
step 1 using
long reads
Illumina reads
200X
iterative by
kmer size
40,50,60
nthit
ntedit
step 2 using
short reads
kmer approach
Illumina reads
200X
x3
pilon
step 3 mapping
short reads
assembly polished
fasta
validations:
- metrics
- BUSCO Analysis
- 100-mer completeness
ONT reads
100X
repeat identification
window masker
purge-haplotigs
genome size
reduction
bionano
Haploid assembly
fasta
validations:
- metrics
- BUSCO Analysis
- 100-mer completeness
HiC data
salsa
Scaffolding
Chr scale assembly
fasta
-genetic map
-sex-linked genes
-transcriptomic female/male data
-female genome data
ALLMAPS mapping
Anchoring the sex
chromosome
caracterization
chromosomes
molecules

**Fig. S1.**

**Genome Assembly Workflow**. This scheme illustrates the comprehensive genome assembly process and each phase. Initially, long reads were assembled using Flye assembler (*43*). Subsequently, a polishing was performed using a multi-step approach, a series of iterations were computed using both long and short reads. To identify and eliminate duplication sequences we use purge haplotigs software. Scaffolding was done including first Bionano and then Hi-C data. To further enhance scaffolding accuracy and resolution, the scaffolds were anchored to chromosomes based on the genetic map using the ALLMAPS algorithm (*52*). Finally, the construction of chromosome Y involved assembling unplaced scaffolds, coverage ratio discrepancy between male and female, and meticulous manual curation of long range Hi-C data. The present workflow ensured the generation of a high-quality chromosome scale genome assembly.

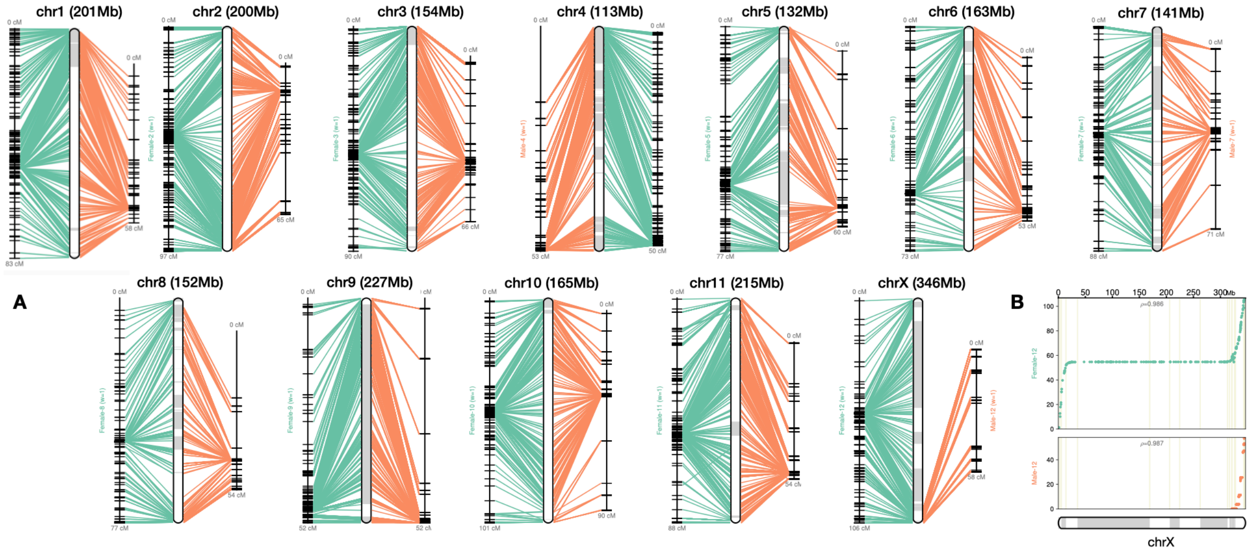


**Fig. S2.**

**Scaffold anchoring onto the genetic map.** A) Visual representation of the anchoring process based on Bionano+Omni-C scaffolds onto the corresponding positions in the genetic map and alignment of the scaffolds along the chromosomes; in green the female markers and in orange the male genetic markers. B) Marey map view of chromosome X in which the x-axis represents the physical positions along the chromosome in Mb, and the y-axis represents the positions on the genetic map in centimorgan (cM).

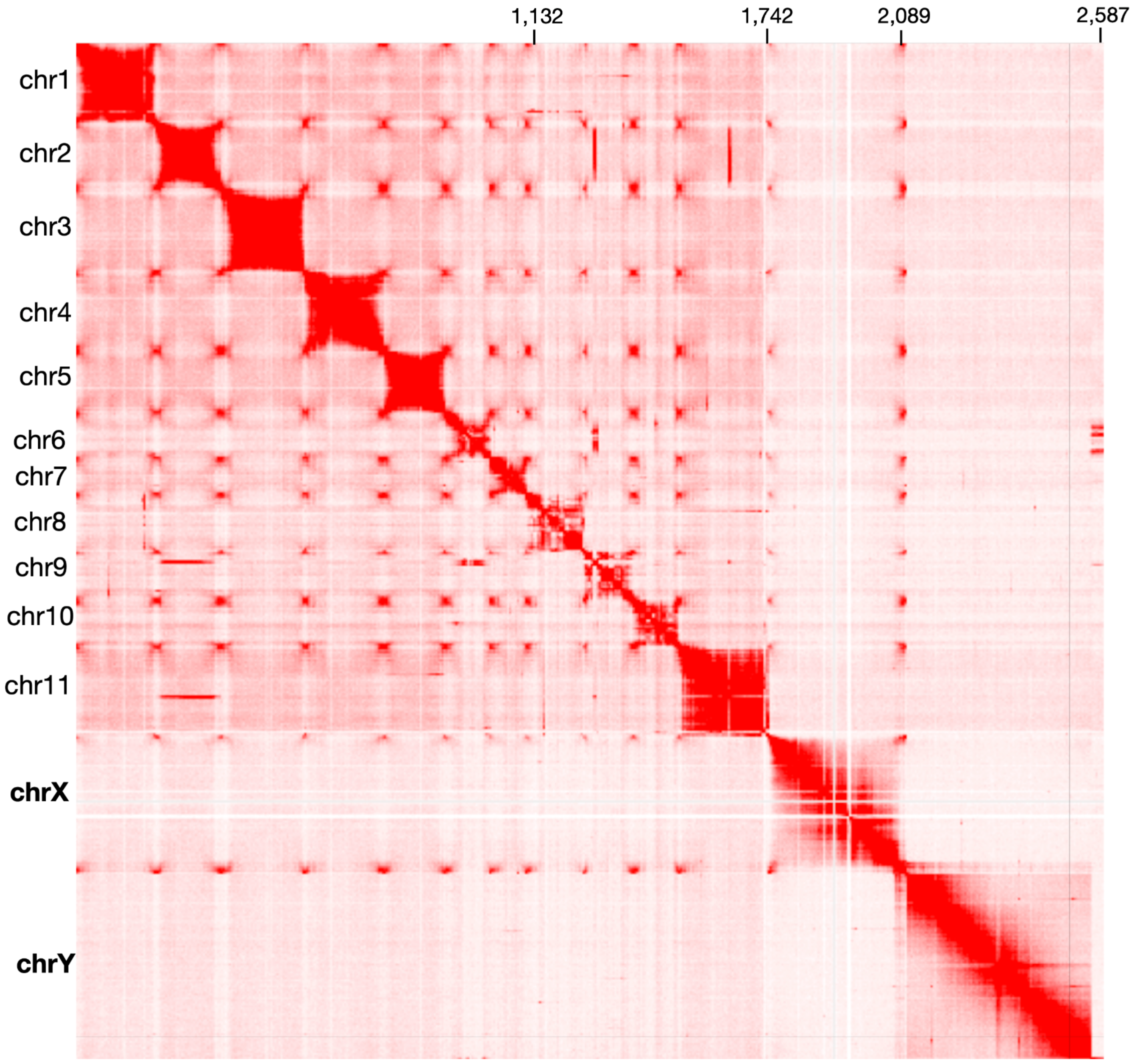


**Fig. S3.**

**Heatmap of Omni-C data.** The heatmap provides a visual representation of the spatial organization of chromatin interaction in the assembly. X and Y axes represent genomic coordinates along the chromosomes. Red color intensity reflects the strength of interactions between the corresponding genomic regions. The diagonal pattern represents the consistency between the chromosome-scale assembly and the long range Hi-C data.

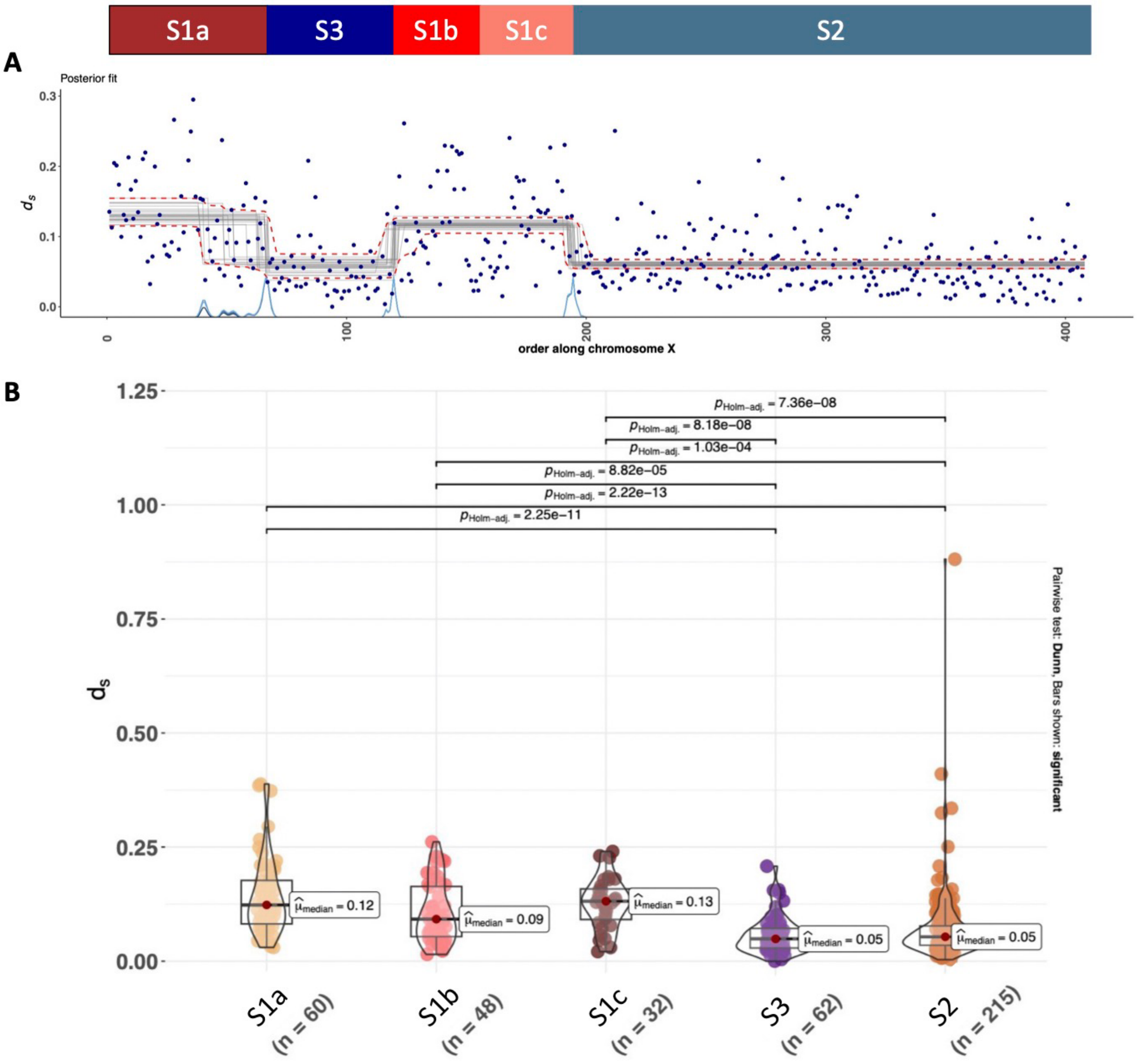


**Fig. S4.**

**Analysis of evolutionary strata.** (A) Changepoint analysis of $d_S$ along the X chromosome. The X-axis shows the gene order on the X. Blue peaks are showing inferred breakpoints. Gray lines show the average dS of the inferred blocks. (B) Statistical test of the median dS among the strata.

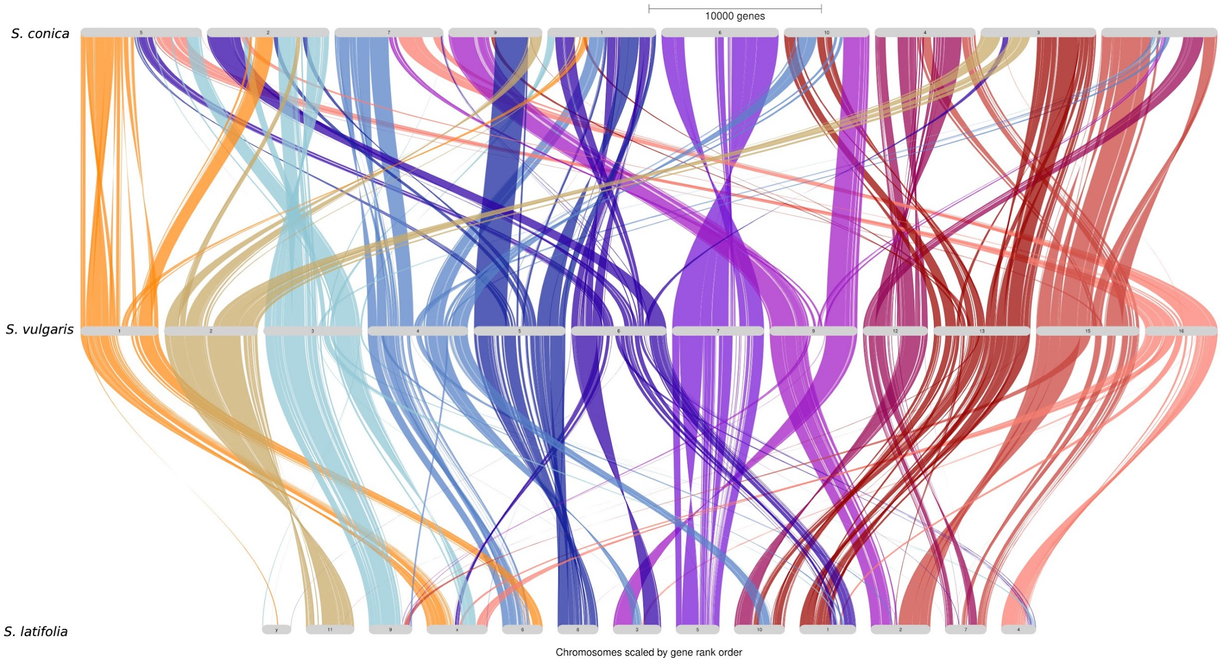


**Fig. S5.**

**Synteny analysis of *S. latifolia, S. conica* and *S. vulgaris* genomes**. Syntenic blocks are shown. Colors are based on *S. vulgaris* chromosomes.

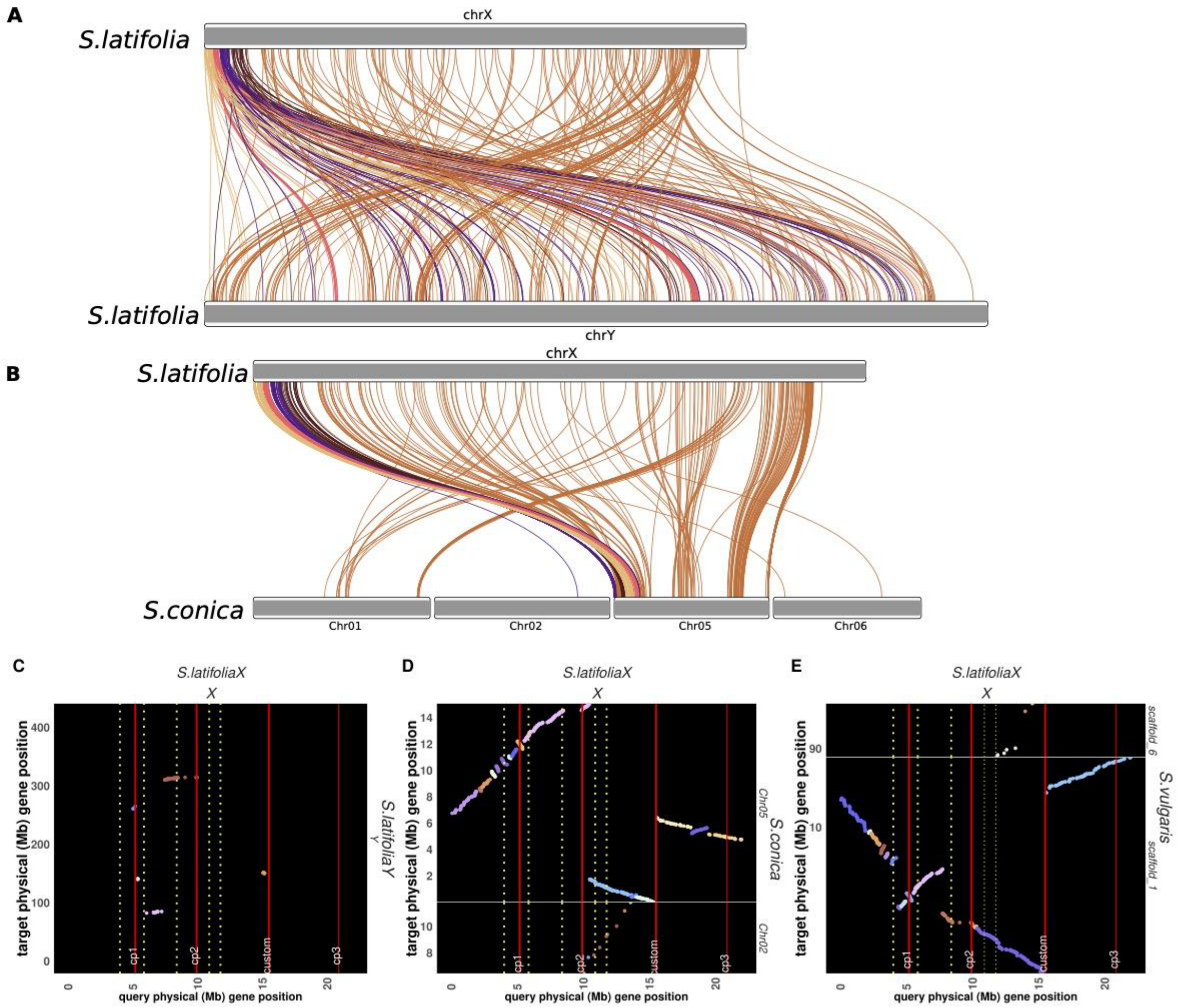


**Fig. S6.**

**Synteny analysis of *S. latifolia* sex chromosomes**. (A) *S. latifolia* X and Y chromosomes comparison. (B) *S. latifolia* X and *S. conica* 1, 2, 5, 6 chromosomes comparison. Dot plots of *S. latifolia* X versus *S. latifolia* Y, (C) *S. latifolia* X versus *S. conica* 5 and 2 (D), *S. latifolia* X versus *S. vulgaris* 6 and 1 (E). All dot plots focus on the first 25 Mb of the *S. latifolia* X chromosome where all strata boundaries are found. The strata boundaries are shown with red vertical lines. cp 1 = stratum 1a/stratum 3 limit, cp2 = stratum 2/stratum 1b limit, custom = stratum 1b/stratum 1c limit, cp 3 = stratum 1c/stratum 2 limit.

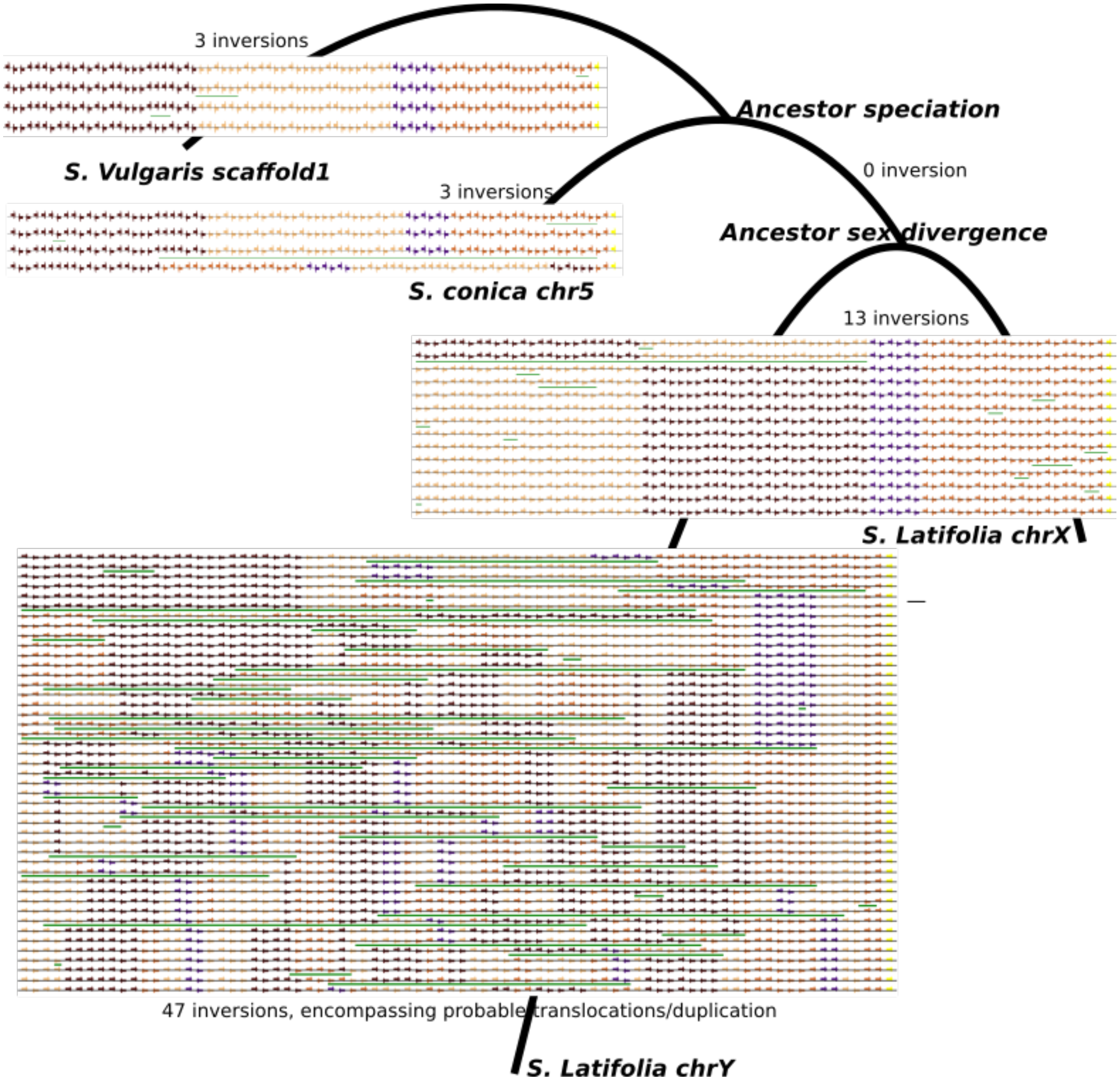


**Fig. S7.**

**Reconstruction of chromosomal rearrangements between the *S. latifolia* X and Y and the outgroups (focus on stratum 1)**. Strata S3 has been removed. Ancestral states were obtained with Badger (*81*), and inversion scenarios with DCJ2HP (*82*). These tools explain all arrangement differences with inversions, which gives an idea of probable ancestral states and of synteny conservation in the different chromosomes, but not every inversion seems plausible. First, because there are a big number of different inversion scenarios to chromosome Y, second because some patterns could be more probably explained by translocations or duplications. Colors of the genes reflect the strata as computed with the changepoint analysis based on sequence divergence. Inversions are shown with green lines. Tree is unrooted.

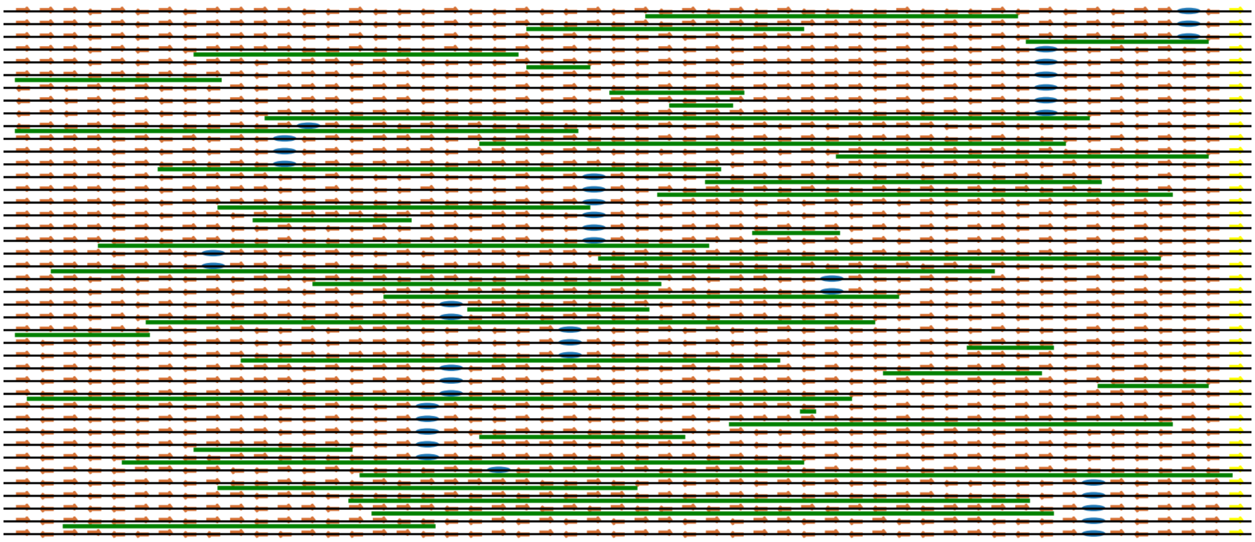

**Fig. S8.**

**Reconstruction of chromosomal rearrangements between the *S. latifolia* X and Y and the outgroups (focus on stratum 2)**. Same legend as figure S8. Centromere of X and its homologous pseudocentromere on the Y (*19*) are depicted by a blue oval.

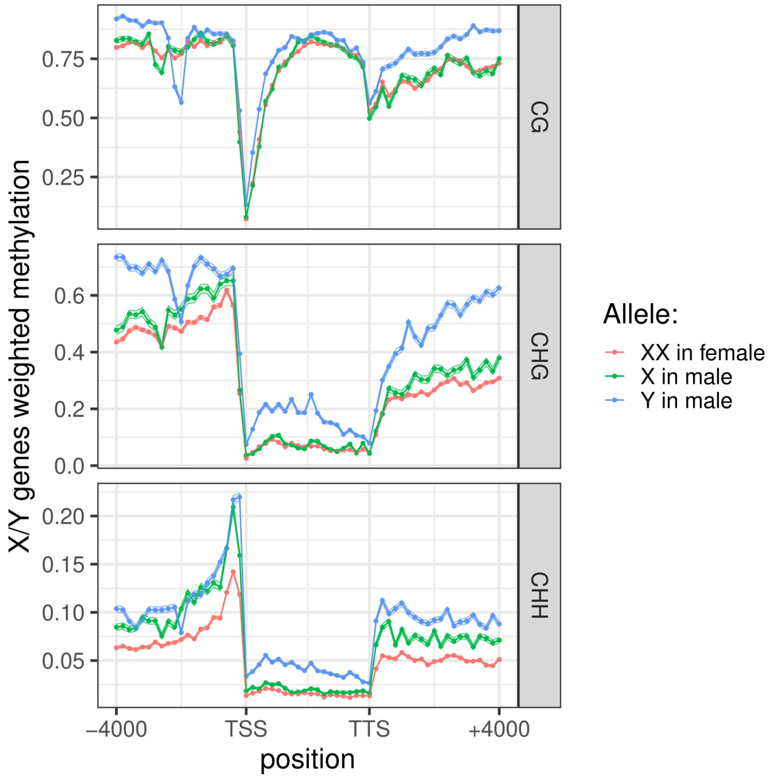


**Fig. S9.**
**Metaplot of X/Y genes DNA methylation in CG (top panel), CHG (middle panel) and CHH context (bottom panel)**. Both X alleles in the female are represented in red, the X allele in the male in green and the Y allele in the male in blue. All X/Y genes were combined to plot the average proportion of methylated reads at cytosine positions along sliding windows; the 95% confidence interval is represented as a small ribbon around the curve. Twenty windows of 200 bp were studied upstream of the transcription start site (TSS) and twenty windows of 200 bp were studied downstream of the transcription termination site (TTS). The gene body (from TSS to TTS) was divided into twenty windows of equal size.

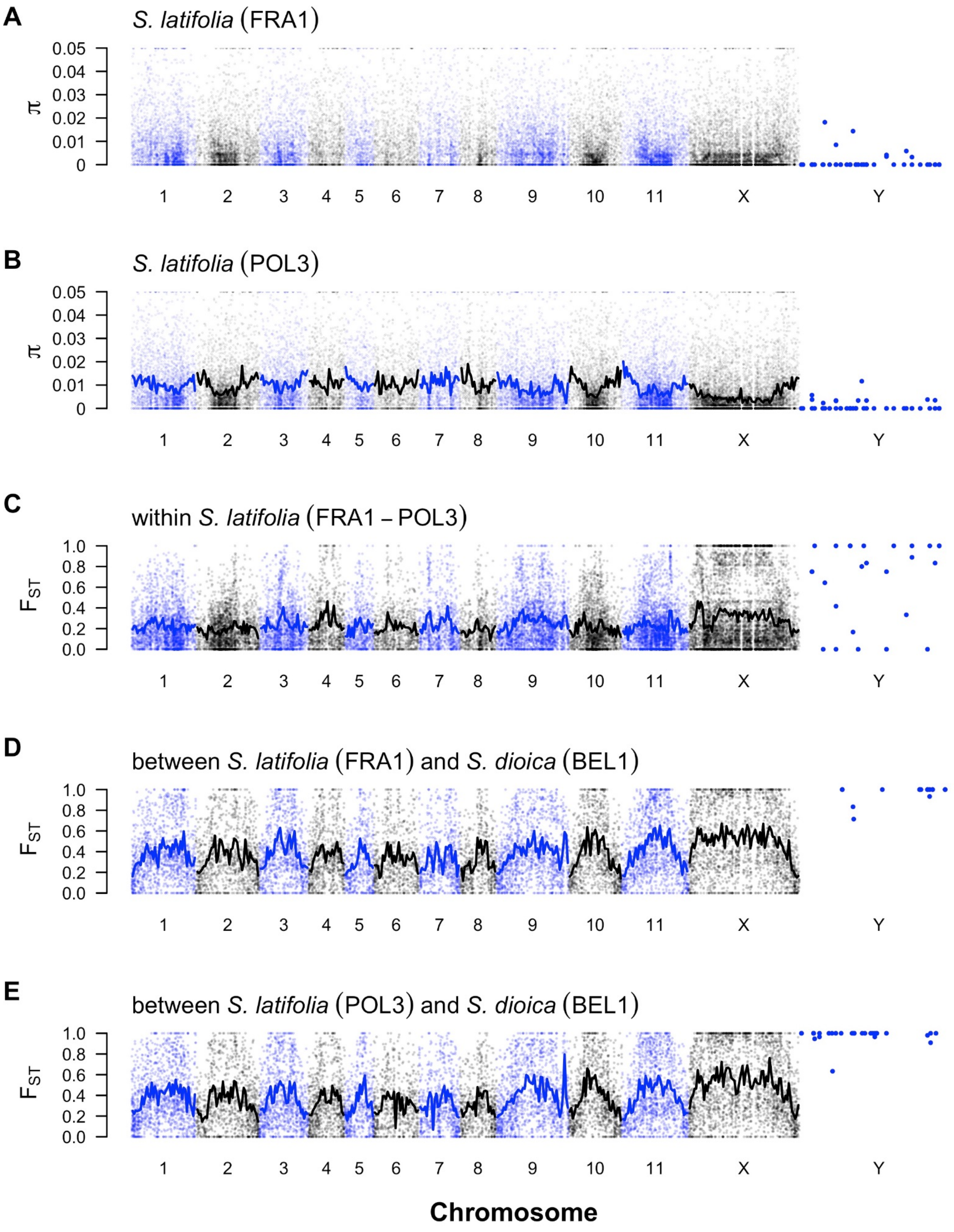


**Fig. S10.**

**Genetic diversity analysis.** Nucleotide diversity $\pi$ (A, B) in two populations of *Silene latifolia* and genetic differentiation, Hudson's $F_{ST}$ (C, D, E) between two populations of *S. latifolia* (C) and between *S. latifolia* and the closely related *S. dioica* (D, E). Given are average values for RADtags (points) with lines based on 5Mb windows along the genome, except for the Y chromosome where data for individual Y-specific RADtags is given. Data for the autosomes and X chromosome were calculated for females, and data for the Y chromosome for males.

A

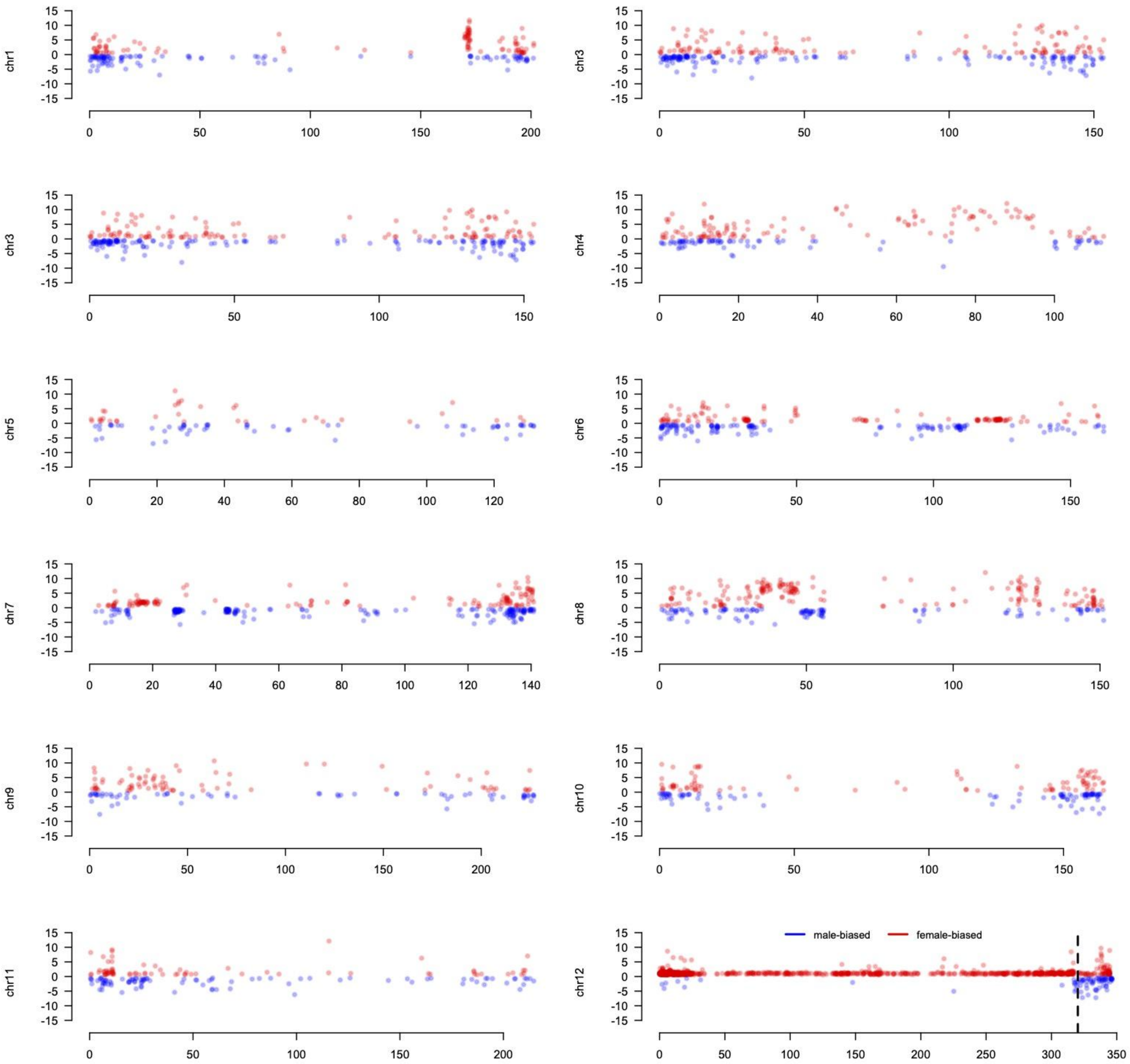

chr1
chr3
chr3
chr4
chr5
chr6
chr7
chr8
chr9
chr10
chr11
chr12
male-biased
female-biased

B

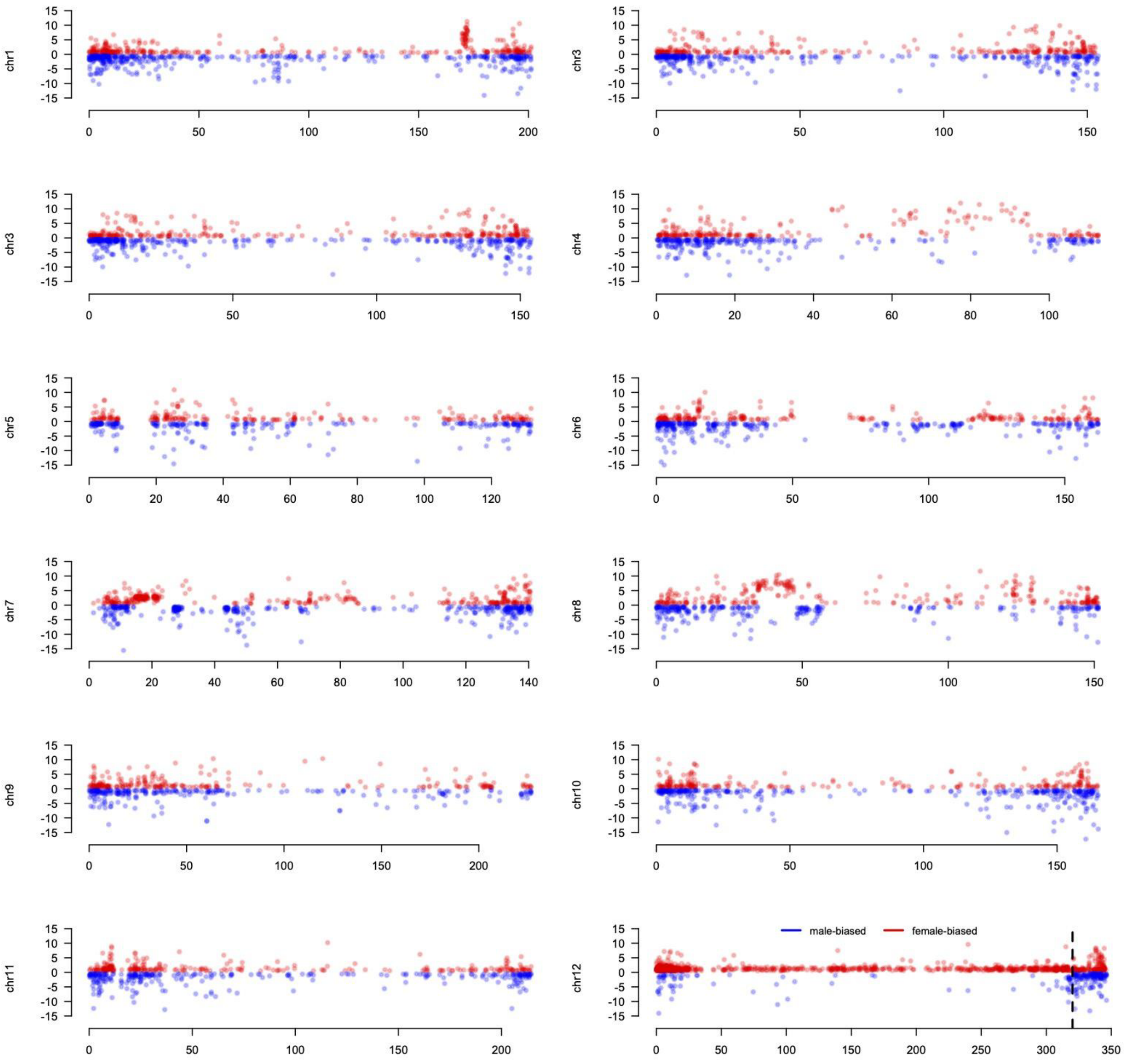


**Fig. S11.**

**Differential expression analysis between male and female flowers at stages 5 and 8 of flower development**. A) for stage 5: 3654 DEGs (1376 down and 2278 up); B) For stage 8: 7928 DEGs (3836 down and 4096 up). Y-axis is showing the log fold change of Female to Male expression, X-axis is showing the position on the chromosomes. Chr12 is the X chromosome, the dotted line indicates the pseudoautosomal boundary. Red dots show the female-biased genes; blue dots show the male-biased genes. No DEG analysis was performed for the Y chromosome as it is not expressed in females.

**Table S1.**

**Sequencing and scaffolding data.**

A) Metrics of sequencing data.

| | **ONT**<br>**6 PromethION** | **PCR-Free**<br>**Illumina** | **Hi-C**<br>**Illumina** |
|---|---|---|---|
| **Cumulative size** | 295.4Gb | 599.9 Gb | 65.7 Gb |
| **# of reads** | 17.8 M | 1,988 M<br>2x151bp | 217.7 M<br>2x151bp |

B) Metrics of optical mapping data.

| | **1 BNG Flow cell (molecules > 150Kb)** | **DLE-1 Optical Map** | **DLE-1 Filtered Optical Map** |
|---|---|---|---|
| **Cumulative size** | 512 Gbp | 4.5 Mb | 2.29 Mb |
| **# of genome maps** | | 944 | 744 |
| **N50** | 207.3 Kb | 20.58 Mb | 16.7 Mb |

C) Metrics of Hybrid scaffolds.

| **# scaffolds** | **1,274** |
|---|---|
| **Cumulative size** | 2,848,429,323 |
| **N50**<br>**(L50)** | 45,710,048<br>(17) |
| **N90**<br>**(L90)** | 3,154,060<br>(114) |
| **Average size** | 2,235,816 |
| **Max size** | 170,686,516 |

**Table S2.**

**Detailed assembly metrics.** Assembly metrics describe molecules as contigs, scaffolds or chromosomes; Total size of the sequences and N50, N90 values, and QV computed using high quality Illumina reads with the Yak software. For Flye we reported QV values before and after the polishing step. BUSCO metrics are reported for all the assemblies using the viridiplantae database (*47*).

| | | **Flye** | **Flye-Purge** | **Bionano** | **Hi-C** | **Allmaps** |
|---|---|---|---|---|---|---|
| **Assembly metrics** | Molecules | contigs | contigs | scaffolds | scaffolds | chromosomes |
| | Number | 6,608 | 1,553 | 1,274 | 912 | 719 |
| | Total Size | 3,746,475,938 | 2,709,641,343 | 2,848,335,899 | 2,848,530,399 | 2,739,705,433 |
| | Longest object | 86,839,985 | 87,054,625 | 170,681,572 | 364,495,766 | 497,814,031 |
| | N50 | 8,692,806 | 16,708,860 | 45,707,540 | 124,369,005 | 200,709,468 |
| | N90 | 315,568 | 2,042,593 | 3,153,934 | 5,712,354 | 112,764,170 |
| | QV | 19.673/29.387 | 29.76 | 29.745 | 29.719 | 29.716 |
| **BUSCO** | Complete % | 99.5 | 92.5 | 92.2 | 92.2 | 92.3 |
| | Complete and single-copy % | 55.5 | 77.2 | 76.9 | 76.9 | 77.2 |
| | Complete and duplicated % | 44 | 15.3 | 15.3 | 15.3 | 15.1 |
| | Fragmented % | 0.2 | 0.9 | 0.9 | 0.9 | 0.9 |
| | Missing % | 0.3 | 6.6 | 6.9 | 6.9 | 6.8 |
| | Total groups searched | 425 | 425 | 425 | 425 | 425 |

**Table S3.**

**Repeat annotation.** The table represents all the annotated principal repeat families (i.e. 61.18 out of 79.24 % total masked repeats).

| | Repeat annotation | Number | Length(bp) | % of repeats | % of the genome |
|---|---|---|---|---|---|
| **LTR** | Ty1/Copia | 25800 | 528,218,684 | 33.88% | 20.73% |
| | Ty3/Gypsy | 23220 | 858,707,731 | 55.07% | 33.69% |
| **DNA** | hAT | 2859 | 3,134,946 | 0.20% | 0.12% |
| | CACTA | 4197 | 10,428,876 | 0.67% | 0.41% |
| | PIF/Harbinger | 436 | 892,063 | 0.06% | 0.04% |
| | Mutator | 3720 | 8,336,613 | 0.53% | 0.33% |
| | Tcl/Mariner | 1063 | 2,692,989 | 0.17% | 0.11% |
| | Helitron | 224330 | 79,203,766 | 5.08% | 3.11% |
| **MITE** | hAT | 3305 | 1,485,132 | 0.10% | 0.06% |
| | CACTA | 86 | 28,482 | <0.01% | <0.01% |
| | PIF/Harbinger | 1084 | 365,176 | 0.02% | 0.01% |
| | Mutator | 1607 | 531,462 | 0.03% | 0.02% |
| | Tcl/Mariner | 4415 | 1,042,309 | 0.07% | 0.04% |
| **LINE** | LINE | 7740 | 12,109,477 | 0.78% | 0.48% |
| **Satellite** | Satellite | 12900 | 52,090,963 | 3.34% | 2.04% |
| **Total repeats** | **-** | - | 1,559,268,669 | 100.00% | 61.18% |

**Table S4.**

**Mutants data**. Samples whose Illumina reads were mapped to the masked reference male genome to detect deleted regions and genes within. First column identifies the sample ID used throughout the manuscript (technical replicates are identified with A and B); second column identifies the individual; third and fourth column label the phenotype of the individual – hermaphrodite (⚥) and asexual with early, intermediate and late arrest in the anther development – and the expected deleted region – GSF, SPF, or MFF); the fifth to the eighth column show the number of reads in each step and derived statistics (specifically, the number of reads obtained from sequencing and the sequence depth, followed by the number of reads kept after the FASTP filtering (*57*), the number of reads that mapped to the reference male genome, and the subset number of reads with a mapping quality (MAPQ) greater or equal to 30). Note that reads for typical males and females are a subset of the total number of reads obtained in this study.

| **Sample ID** | **Indivi-dual ID** | **Phenotype** | **Dele-ted region** | **Initial no. of reads** | **Sequence depth** | **No. of reads after FASTP filter** | **No. of mapped reads onto the masked genome** | **No. of mapped reads after filtering MAPQ ≥ 30** |
|---|---|---|---|---|---|---|---|---|
| CM | - | Typical male | - | 397,691,268 | 21X | 356,025,000 | 352,602,551 | 237,936,765 |
| CF | - | Typical female | - | 158,674,092 | 8X | 156,560,864 | 154,871,047 | 87,385,638 |
| Ind1.A | MH14 | ⚥ | GSF | 187,836,438 | 10X | 181,935,544 | 142,385,401 | 71,348,204 |
| Ind1.B | MH14 | ⚥ | GSF | 432,121,224 | 23X | 106,122,902 | 101,227,775 | 45,400,827 |
| Ind2.A | MH115 | ⚥ | GSF | 116,013,362 | 6X | 111,234,108 | 109,411,682 | 57,142,213 |
| Ind2.B | MH115 | ⚥ | GSF | 49,036,446 | 2X | 37,298,956 | 33,357,584 | 15,689,074 |
| Ind3.A | MH12 | ⚥ | GSF | 140,619,190 | 7X | 137,617,504 | 136,114,346 | 71,707,755 |
| Ind3.B | MH12 | ⚥ | GSF | 120,859,654 | 6X | 202,894,040 | 200,386,646 | 101,992,306 |
| Ind4.A | UH17 | ⚥ | GSF | 94,110,610 | 5X | 91,570,138 | 90,493,599 | 41,882,897 |
| Ind4.B | UH17 | ⚥ | GSF | 50,645,318 | 2X | 36,708,438 | 31,101,982 | 11,459,495 |
| Ind5.A | MH5 | ⚥ | GSF | 65,548,068 | 3X | 64,273,754 | 63,646,537 | 31,945,547 |
| Ind5.B | MH5 | ⚥ | GSF | 217,595,746 | 11X | 206,109,490 | 201,905,256 | 97,291,913 |
| Ind6 | MH78 | ⚥ | GSF | 297,753,556 | 16X | 203,374,404 | 194,810,831 | 93,075,849 |
| Ind7 | MH79 | ⚥ | GSF | 368,457,918 | 20X | 298,440,444 | 271,861,153 | 120,916,254 |

| | | | | | | | | |
|---|---|---|---|---|---|---|---|---|
| Ind8 | UH13 | ⚥ | GSF | 337,683,810 | 18X | 288,143,016 | 270,533,488 | 117,305,070 |
| Ind9 | UH15 | ⚥ | GSF | 349,365,674 | 19X | 230,628,346 | 222,100,229 | 99,437,922 |
| Ind10.A | MS37 | Early arrest | SPF | 141,505,176 | 7X | 138,866,048 | 137,411,545 | 69,666,321 |
| Ind10.B | MS37 | Early arrest | SPF | 151,056,598 | 8X | 132,538,968 | 126,008,879 | 60,114,872 |
| Ind11 | MS36 | Early arrest | SPF | 389,708,634 | 21X | 322,678,162 | 298,080,365 | 129,116,005 |
| Ind12.A | MS34 | Intermediate arrest | SPF | 33,933,558 | 1X | 33,035,306 | 32,697,773 | 16,367,876 |
| Ind12.B | MS34 | Intermediate arrest | SPF | 234,192,048 | 12X | 38,652,335 | 37,111,988 | 17,520,604 |
| Ind13.A | MS65 | Intermediate arrest | SPF | 117,143,684 | 6X | 114,613,254 | 113,353,261 | 55,348,621 |
| Ind13.B | MS65 | Intermediate arrest | SPF | 220,181,036 | 12X | 187,704,346 | 174,814,748 | 77,803,261 |
| Ind14 | MS75 | Intermediate arrest | SPF | 369,561,884 | 20X | 333,011,070 | 320,025,818 | 146,664,950 |
| Ind15.A | MS96 | Late arrest | MFF | 122,311,938 | 6X | 120,101,614 | 118,860,836 | 61,741,114 |
| Ind15.B | MS96 | Late arrest | MFF | 257,082,764 | 14X | 146,762,492 | 143,148,780 | 70,583,897 |
| Ind16 | MS62 | Late arrest | MFF | 314,478,774 | 17X | 265526898 | 247281567 | 85,280,811 |
| Ind17 | US39 | Late arrest | MFF | 189,263,204 | 10X | 55,656,126 | 50,014,792 | 18,985,743 |
| Ind18 | MS63 | Late arrest | MFF | 136,974,320 | 7X | 134,379,628 | 132,896,022 | 68,069,072 |

**Table S5.**
**Transcriptomic data**. Flower sampling for RNA-seq analysis and number of read counts.

| Sexual morph | Flower stage | Samples | Read counts (millions) | Normalized read counts (millions) |
|---|---|---|---|---|
| Male | Stage 5 | F_S5_R12 | 17.837768 | 17.326070 |
| | | F_S5_R13 | 19.535579 | 17.786076 |
| | | F_S5_R14 | 17.172442 | 17.350641 |
| | Stage 8 | F_S8_R15 | 18.244499 | 17.050744 |
| | | F_S8_R17 | 16.96041 | 17.143541 |
| | | F_S8_R19 | 17.534571 | 18.014962 |
| Female | Stage 5 | M_S5_R5 | 17.670883 | 17.145611 |
| | | M_S5_R8 | 17.25855 | 17.124977 |
| | | M_S5_R9 | 17.02373 | 16.772271 |
| | Stage 8 | M_S8_R1 | 18.389832 | 18.437642 |
| | | M_S8_R2 | 16.919949 | 19.309282 |
| | | M_S8_R4 | 19.186526 | 18.445721 |
